# Host and viral determinants of airborne transmission of SARS-CoV-2 in the Syrian hamster

**DOI:** 10.1101/2022.08.15.504010

**Authors:** Julia R. Port, Dylan H. Morris, Jade C. Riopelle, Claude Kwe Yinda, Victoria A. Avanzato, Myndi G. Holbrook, Trenton Bushmaker, Jonathan E. Schulz, Taylor A. Saturday, Kent Barbian, Colin A. Russell, Rose Perry-Gottschalk, Carl I. Shaia, Craig Martens, James O. Lloyd-Smith, Robert J. Fischer, Vincent J. Munster

## Abstract

It remains poorly understood how SARS-CoV-2 infection influences the physiological host factors important for aerosol transmission. We assessed breathing pattern, exhaled droplets, and infectious virus after infection with Alpha and Delta variants of concern (VOC) in the Syrian hamster. Both VOCs displayed a confined window of detectable airborne virus (24-48 h), shorter than compared to oropharyngeal swabs. The loss of airborne shedding was linked to airway constriction resulting in a decrease of fine aerosols (1-10µm) produced, which are suspected to be the major driver of airborne transmission. Male sex was associated with increased viral replication and virus shedding in the air. Next, we compared the transmission efficiency of both variants and found no significant differences. Transmission efficiency varied mostly among donors, 0-100% (including a superspreading event), and aerosol transmission over multiple chain links was representative of natural heterogeneity of exposure dose and downstream viral kinetics. Co-infection with VOCs only occurred when both viruses were shed by the same donor during an increased exposure timeframe (24-48 h). This highlights that assessment of host and virus factors resulting in a differential exhaled particle profile is critical for understanding airborne transmission.

## Introduction

Transmission by aerosolized virus particles has been a major contributor to the spread of SARS-CoV-2 [1, 2] [3–6]. Although highly efficient in preventing severe disease, vaccines do not significantly reduce transmission of variants of concern (VOCs) [7]. Transmission occurs when people release respiratory droplets carrying virus during (e.g.) speaking, singing, breathing, sneezing, or coughing. Droplet size and half-life in the air are not uniform [8, 9] and depend on speech and breathing patterns [10], COVID-19 severity, and physiological parameters such as age [11, 12]. As with influenza [13], SARS-CoV-2 RNA was detectable mostly in fine aerosols in humans, as opposed to coarse aerosols [11]. It is not clear how exhaled droplet size, breathing patterns and even the quantity of exhaled infectious virus itself fundamentally contribute to the airborne transmission efficiency *in vivo* and how COVID-19 directly influences additional physiological factors which may contribute to fine aerosol production. There is reportedly large heterogeneity in the transmission potential of individuals. Superspreading events have been reported numerous times throughout the pandemic and are suggested to be a major driver [14, 15]. They are thought to arise from a combination of biological, social, and chance factors. While human epidemiology and modeling studies have highlighted various factors which may contribute to SARS-CoV-2 transmission heterogeneity, including viral load [16], much of the observed variance remains poorly understood. These factors are currently best studied in small animal models like the Syrian hamster, which allow for stringent and controlled experimental comparisons. The Syrian hamster model has been widely used to study SARS-CoV-2 transmission [17]; it recapitulates human contact, fomite and, importantly, airborne short distance and fine aerosol transmission [2, 18–20]. In this model, highest efficiency of short-distance airborne transmission was observed before onset of weight loss and acute lung pathology, peaking at one day post inoculation and correlating to the highest virus loads in the upper respiratory tract of donor animals [21]. Data on lung function loss in the Syrian hamster model after SARS-CoV-2 infection is available [22, 23], and virus has been demonstrated in exhaled droplets [24]. Yet, a systematic study that addresses how airborne transmission potential depends on these features, along with recognized influences of sex and VOC, has not been performed. The study of these contributing factors would allow us to address how they come together to shape transmission outcomes.

Here, we introduce a mathematical model delineating for Alpha and Delta VOCs the relationship between exhaled infectious virus and virus detected in the upper respiratory tract during infection and longitudinally detail the changes in lung function, respiratory capacity, and exhaled particle profiles. Finally, we assess the airborne transmission competitiveness and heterogeneity *in vivo* of Alpha and Delta.

## Results

### Peak infectious SARS-CoV-2 in air samples is detected between 24 h and 48 h post infection

Structural modelling and pseudotype-entry comparison suggested that the Syrian hamster model should recapitulate the entry-specific competitive advantage of Delta over Alpha observed in humans (**Figure S 1A-D).** Syrian hamsters were inoculated with a low dose (10^3^ TCID_50_, intranasal (IN), N = 10 per group) of SARS-CoV-2 Delta or Alpha. Animals were monitored for 14 days post inoculation (DPI). We observed no significant differences in weight loss or viral titers in lung or nasal turbinates between the variants (**Figure S 2A-C**). At 14 DPI, hamsters (N = 5) mounted a robust anti-spike IgG antibody response, and the overall binding pattern was similar between Alpha and Delta (**Figure S 2D,E**). In a live virus neutralization assay, homologous virus was neutralized significantly better as compared to the heterologous variant (**Figure S 2F,G**), but no significant difference was determined between the neutralization capacity against the respective homologous variant (median reciprocal virus neutralization titer = 320 (Alpha anti-Alpha)/ 320 (Delta anti-Delta), p = 0.9568, N = 5, ordinary two-way ANOVA, followed by Tukey’s multiple comparisons test).

We determined the window of SARS-CoV-2 shedding for Alpha and Delta using swabs from the upper respiratory tract and air sampling from cages, quantifying virus using gRNA, sgRNA, and infectious virus titers. Oral swabs remained positive for gRNA and sgRNA until 7 DPI, but infectious virus dropped to undetectable levels after 4 DPI in most individuals (**Figure S 3A)**. Cage air was sampled during the first 5 days of infection in 24 h time windows from cages containing 2 or 3 animals, grouped by sex. gRNA and sgRNA were detectable as early as 1 DPI in 50% of air samples and remained high through 5 DPI, while infectious virus peaked at on 2 DPI and was detectable for a shorter window, from 1 to 4 DPI (**Figure S 3B)**.

### Mathematical modeling demonstrates airborne shedding peaks later and declines faster than oral swab viral load

We quantified heterogeneity in shedding by variant, sex, and sampling method by fitting a mathematical model of within-hamster virus kinetics (see **SI Mathematical Model**) to the data. This served to correlate parameters which are easier to measure, such as RNA in the oral cavity, to the quantity of greatest interest for understanding transmission (i.e. infectious virus in the air per unit time). To do this, we jointly inferred the kinetics of shed airborne virus and parameters relating observable quantities (e.g., plaques from purified air sample filters) to the actual longitudinal shedding. The inferential model uses mechanistic descriptions of deposition of infectious virus into the air, uptake from the air, and loss of infectious virus in the environment to extract estimates of the key parameters describing viral kinetics, as well as the resultant airborne shedding, for each animal. Virus was detectable and peaked earlier in oral swabs (approximately 24 h post inoculation) than virus sampled from the air (approximately 48 h post inoculation), and quantity of detected virus declined slower in the swabs (**Figure 1A,B**). gRNA and sgRNA declined slower than infectious virus both in the air and in swabs. Oral swab data was an imperfect proxy for airborne shedding, even when we directly quantified infectious virus titers. This was due to a lag between peak swab shedding and peak airborne shedding. Inferred within-host exponential growth and decay rates were similar for the two variants. For both variants, males shed more virus than females, even after accounting for males’ higher respiration rates in measurements of shedding into the air. We found a slightly higher ratio of infectious virus to sgRNA in air samples for Delta than for Alpha (**Figure 1B, Figure S 3C,D).** We also found substantial individual-level heterogeneity in airborne shedding, even after accounting for sex and variant (**Figure 1B**). For example, air samples from cage 5 had more than twice as many peak plaques per capita than cage 6, even though both cages contained hamsters of the same sex, inoculated by the same dose, route, and variant. Our model captures this, with substantial inferred heterogeneity in individual airborne shedding in PFU per h, both in timing and in height of peak (**Figure 1B**)

**Figure 1.**
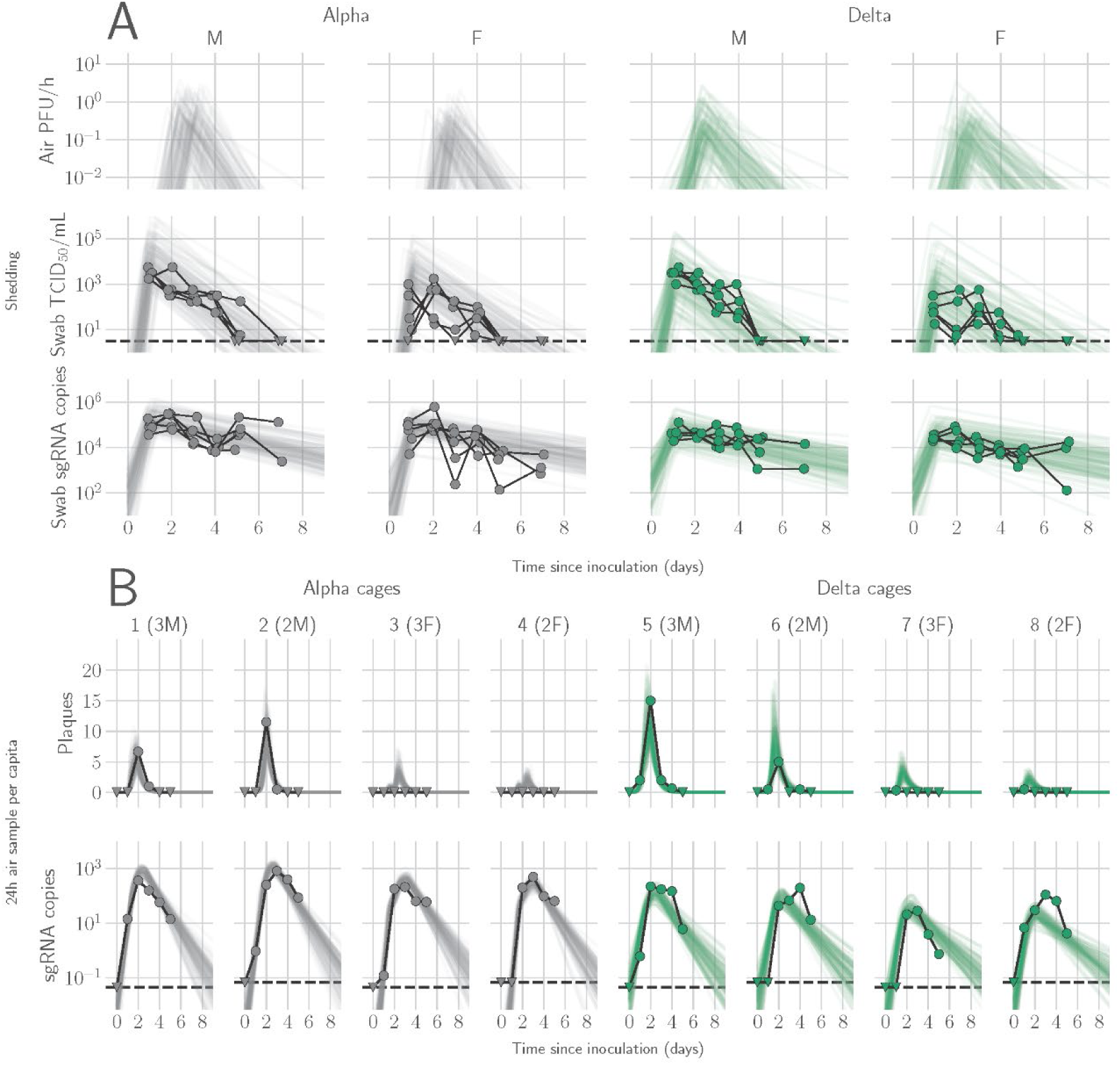
Alpha and Delta variant shedding profiles in oral swabs and air samples. Syrian hamsters were inoculated with 10^3^ TCID_50_ via the intranasal route with Alpha or Delta. **A.** Comparison of swab viral load and virus shedding into the air. Inferred profile of air shedding in PFU/h compared to sgRNA levels and infectious virus titers (TCID_50_/mL) in oropharyngeal swabs collected 1, 2, 3, 4, 5, and 7 DPI. Semitransparent lines are 100 random draws from the inferred posterior distribution of hamster within-host kinetics for each of the metrics. Joined points are individual measured timeseries for experimentally infected hamsters; each set of joined points is one individual. Measurements and inferences shown grouped by variant and animal sex. Measurement points are randomly jittered slightly along the x (time) axis to avoid overplotting. **B**. Viral sgRNA and infectious virus (PFU) recovered from cage air sample filters over a 24 h period starting at 0, 1, 2, 3, 4, and 5 DPI. Points are measured values, normalized by the number of hamsters in the cage (2 or 3) to give per-capita values. Downward-pointing arrows represent virus below the limit of detection (0 observed plaques or estimated copy number corresponding to Ct ≥ 40). Semitransparent lines are posterior predictions for the sample that would have been collected if sampling started at that timepoint; these reflect the inferred underlying concentrations of sgRNA and infectious virus in the cage air at each timepoint and are calculated from the inferred infection kinetics for each of the hamsters housed within the cage. 100 random posterior draws shown for each cage. Cages housed 2 or 3 hamsters; all hamsters within a cage were of the same sex and infected with the same variant. Column titles show cage number and variant, with number of and sex of individuals in parentheses. Dotted lines limit of detection. Grey = Alpha, teal = Delta, p-values are indicated where significant. Abbreviations: sg, subgenomic; TCID, Tissue Culture Infectious Dose; PFU, plaque forming unit; F, female; M, male; DPI, days post inoculation.

### Changes in breathing profile after SARS-CoV-2 infection precede onset of weight loss and are variant and sex-dependent

Pathology in nasal turbinates and lungs did not differ significantly between animals (**Figure S 4**). Pathological changes were consistent with those described previously for COVID-19 in Syrian hamsters after intranasal inoculation with other SARS-CoV-2 strains [19]. Whole body plethysmography (WBP) was performed. We focused the analysis on the first 5 days after inoculation, in which changes in virus shedding and release into the air were observed (**Figure S 2A**). Expiratory time (Te), inspiratory time (Ti), percentage of breath occupied by the transition from inspiration to expiration (TB), end expiratory pause (EEP), breathing frequency (f), peak inspiratory flow (PIFb), peak expiratory flow (PEFb), tidal volume (TVb), minute volume (MVb) and enhanced pause (Penh) were used to assess changes in pulmonary function throughout infection. Principal component analysis was used to determine trends in lung function changes across all groups (**Figure 2A**). This revealed a large degree of inherent variation in individual hamster plethysmography measures. Before inoculation there was no discernible pattern to the clustering observed besides a slight separation by sex. Beginning at 2 DPI, we observed a separation of infected and control animals. This coincided with the observation that all SARS-CoV-2 animals visibly decreased activity levels after 2 DPI, reducing exploratory activity and grooming with sporadic short convulsions which may represent coughing. No single parameter had an overwhelming influence on clustering, though several parameters contributed strongly across all days: Te, Ti, TB, EEP, f, PIFb, PEFb, TVb, and MVb (**Figure 2B,C**).

**Figure 2.**
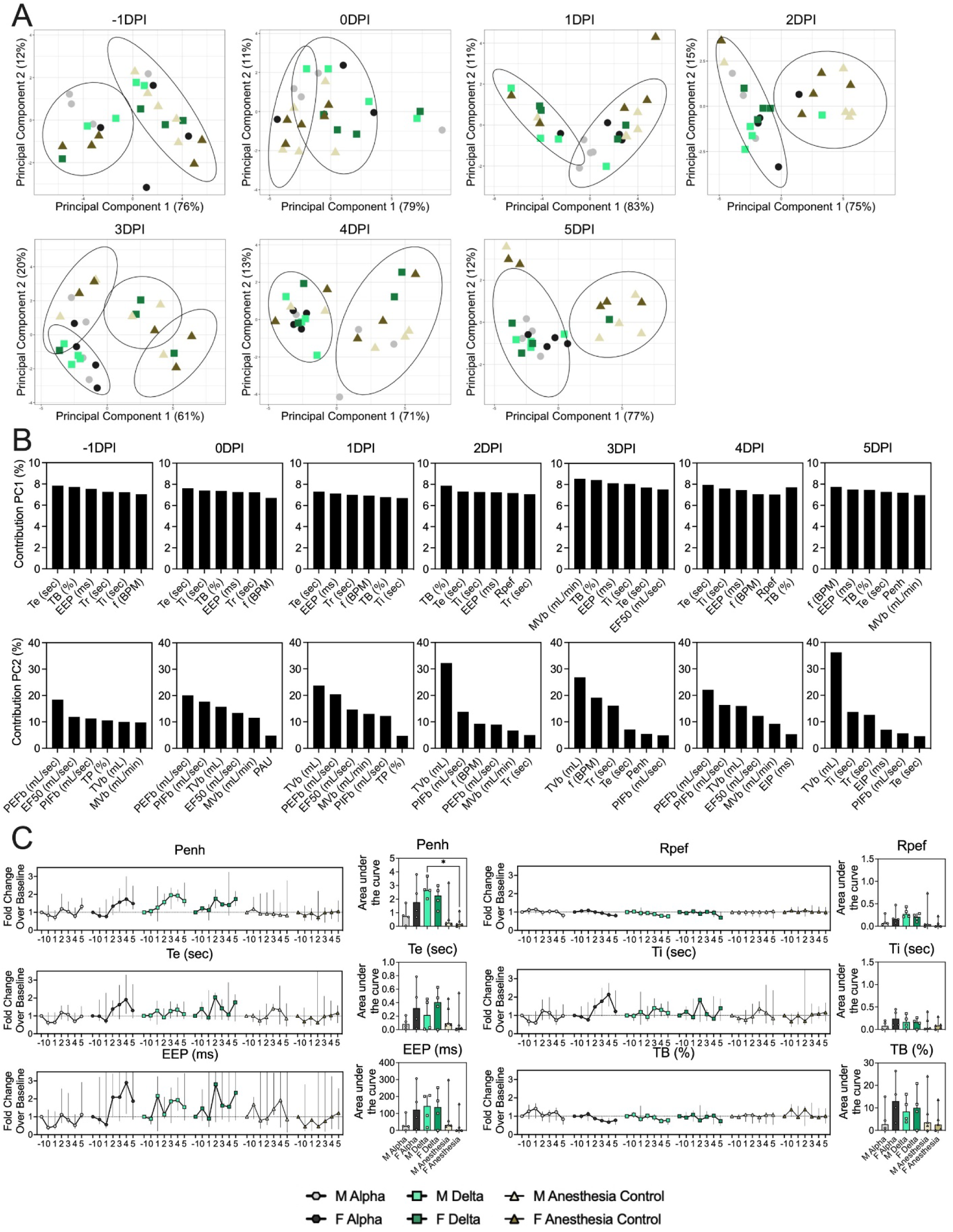
Lung function and breathing changes after SARS-CoV-2 infection with Alpha and Delta. Syrian hamsters were inoculated with 10^3^ TCID_50_ via the intranasal route with Alpha or Delta. **A.** Lung function was assessed on day −1, 0, 1, 2, 3, 4, and 5 by whole body plethysmography. Principal component analysis was used to investigate individual variance. Depicted are principal component (PC) 1 and 2 for each day, showing individual animals (colors refer to legend on right, sex-separated) and clusters (black ellipses). **B.** Individual loading plots for contributions of top 6 variables to PC1 and 2 at each day. **C.** Relevant subset of lung function parameters. Line graphs depicting median and 95% CI fold change values (left) and area under the curve (AUC, right), Kruskal-Wallis test, p-values indicated where significant. Grey = Alpha, teal = Delta, beige = anesthesia control, light = male, dark = female. Abbreviations: Expiratory time (Te), inspiratory time (Ti), percentage of breath occupied by the transition from inspiration to expiration (TB), end expiratory pause (EEP), breathing frequency (f), peak inspiratory flow (PIFb), peak expiratory flow (PEFb), tidal volume (TVb), minute volume (MVb), enhanced pause (Penh), male (M), female (F).

Broad patterns emerged by variant and by sex. Cumulative Penh AUC values for all infected groups were increased compared to the sex-matched control hamsters (p = 0.022, Kruskal-Wallis test, N = 4 for Alpha and Delta, N = 5 for controls). The median Penh AUC values for Alpha, Delta, and control males were 0.741, 2.666, and 0.163, respectively (p = 0.062). The median Penh AUC values for Alpha females, Delta females, and control females were 1.783, 2.255, and 0.159, respectively (p = 0.019). At 4 DPI, the median fold change Penh values for Alpha males and Delta males were 0.793 and 1.929, respectively, as compared to 0.857 for control males. The corresponding Penh values for Alpha, Delta, and control females were 1.736, 1.410, and 1.008, respectively. The separation on 4 DPI did not translate to significant changes in more traditional measures of respiratory function, including f, TVb, and MVb.

### Changes in exhaled aerosol aerodynamic profile after SARS-CoV-2 infection precede acute disease, are variant and sex-dependent

Alpha and Delta inoculated groups (N = 10 each) and a control group (N = 10) were individually evaluated on 0, 1, 3, and 5 DPI. To normalize the particle counts between animals we focused on the percentage of particles in each size range. Across each variant group, particle diameter size <0.53 µm was the most abundant (**Figure 3A**). No consistent, significant overall change in the number of overall particles across all sizes was observed between groups (**Figure S 5C**). Particles between 1 and 10 µm in diameter, most relevant for fine aerosol transmission [25], were examined. At baseline (0 DPI), females across all groups produced a higher proportion of droplets in the 1-10 µm diameter range compared to males (**Figure 3A**). At 3 DPI, the particle profiles shifted towards smaller aerodynamic diameters in the infected groups. At 5 DPI, even control animals demonstrated reduced exploratory behavior, resulting in a reduction of particles in the 1-10 µm range, which could be due to acclimatization to the chamber. This resulted in an overall shift in particle size from the 1-10 µm range to the <0.53 µm range. To analyze these data, individual slopes for each animal were calculated using simple linear regression across the four timepoints (Percent ∼ Intercept + Slope ∗ Day) for percent of particles in the <0.53 µm range and percent of particles in the 1-10 µm range and multiple linear regression was performed (**Figure 3B**). Females had a steeper decline at an average rate of 2.2 per day after inoculation in the percent of 1-10 μm particles (and a steeper incline for <0.53 μm) when compared to males, while holding variant group constant. When we compared variant group while holding sex constant, we found that the Delta group had a steeper decline at an average rate of 5.6 per day in the percent of 1-10 μm particles (and a steeper incline for <0.53 μm); a similar trend, but not as steep, was observed for the Alpha group.

**Figure 3:**
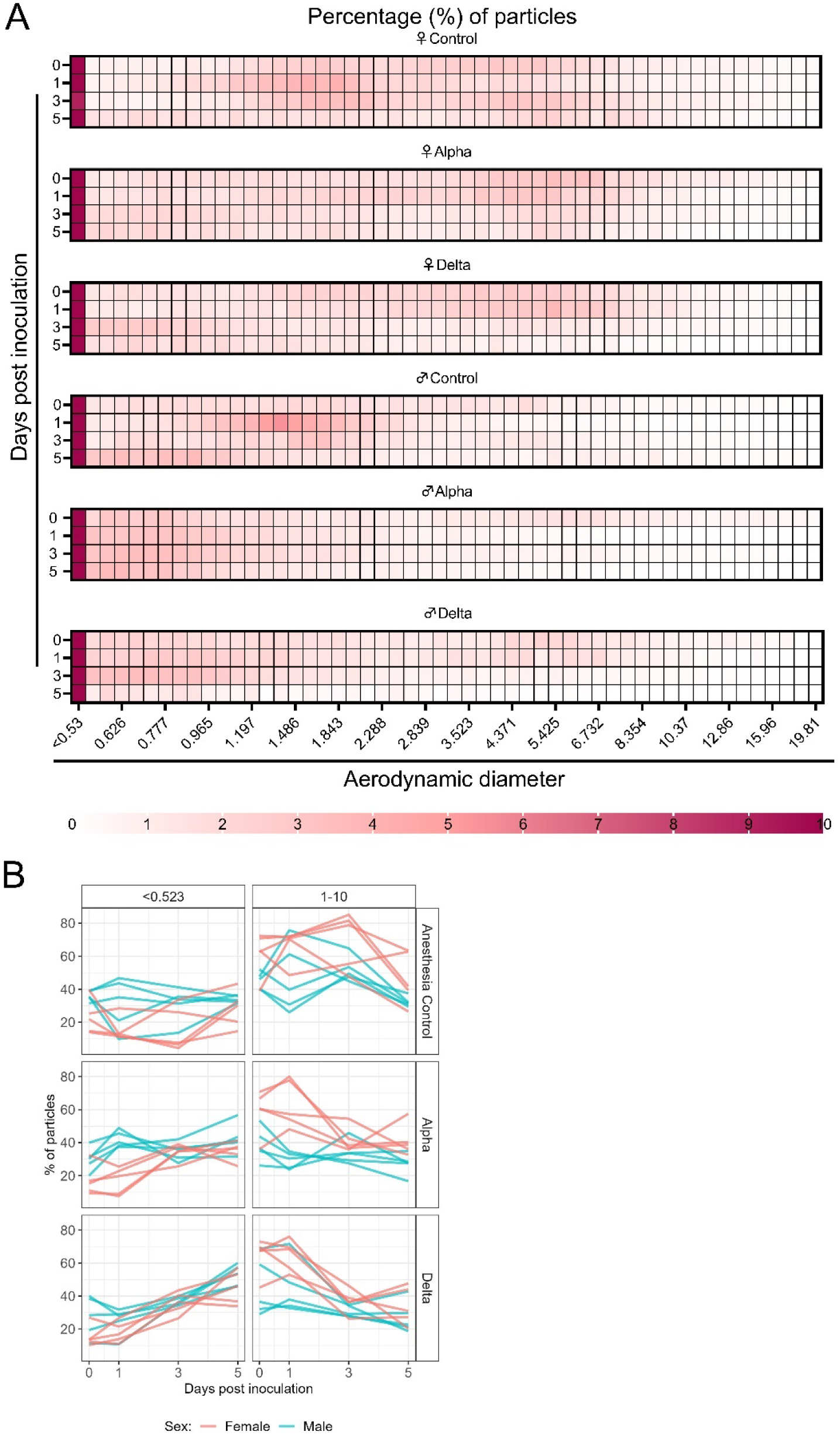
Aerodynamic particle analysis of SARS-CoV-2 infected hamsters. **A.** Syrian hamsters were inoculated with 10^3^ TCID_50_ via the intranasal route with Alpha or Delta. Aerodynamic diameter profile of exhaled particles was analyzed on day 0, 1, 3, and 5. Heatmap shows rounded median percent of total particles across groups, including the anesthesia control group (N = 10, comprising 5 males and 5 females). Colors refer to scale below. **B.** For each animal, line graphs of the percent of particles in the <0.53 and 1-10 µm diameter range by variant group and sex indicated by color. Multiple linear regression performed for each diameter range with group and sex as predictors, F-statistic (3,26) = 9.47 for <0.53 µm model and F-statistic (3,26) = 2.62 for 1-10 µm model, with Tukey multiple comparison adjustment for the three variant-group comparisons (95% family-wise confidence level). For <0.53 range, Male-Female (estimate = −1.7, standard error = 0.888, two-sided p = 0.0659); Alpha-Control (estimate = 2.41, standard error = 1.09, two-sided p = 0.0874), Delta-Control (estimate = 5.40, standard error = 1.09, two-sided p = 0.0001), Delta-Alpha (estimate = 2.99, standard error = 1.09, two-sided p = 0.0280). For 1-10 range, Male-Female (estimate = 2.19, standard error = 1.23, two-sided p = 0.0875); Alpha-Control (estimate = −0.633, standard error = 1.51, two-sided p = 0.9079), Delta-Control (estimate = −3.098, standard error = 1.51, two-sided p = 0.1197), Delta-Alpha (estimate = −2.465, standard error = 1.51, two-sided p = 0.2498). Grey = Alpha, teal = Delta, beige = anesthesia control, red = female, blue = male.

The estimated difference in slopes for Delta vs. controls and Alpha vs. controls in the percent of <0.53 μm particles was 5.4 (two-sided adjusted p= 0.0001) and 2.4 (two-sided adjusted p = 0.0874), respectively. The estimated difference in slopes for percent of 1-10 μm particles was not as pronounced, but similar trends were observed for Delta and Alpha. Additionally, a linear mixed model was considered and produced virtually the same results as the simpler analysis described above; the corresponding linear mixed model estimates were the same and standard errors were similar.

### Alpha and Delta VOC attack rates reveal minimal individual risk of dual infection *in-vivo*

We next compared attack rates between Alpha and Delta during a 4 h exposure window at 200 cm distance. Groups of sentinels (N = 4 or 5) were exposed to two donor animals, one inoculated with Alpha and one inoculated with Delta (**Figure 4A**).

**Figure 4.**
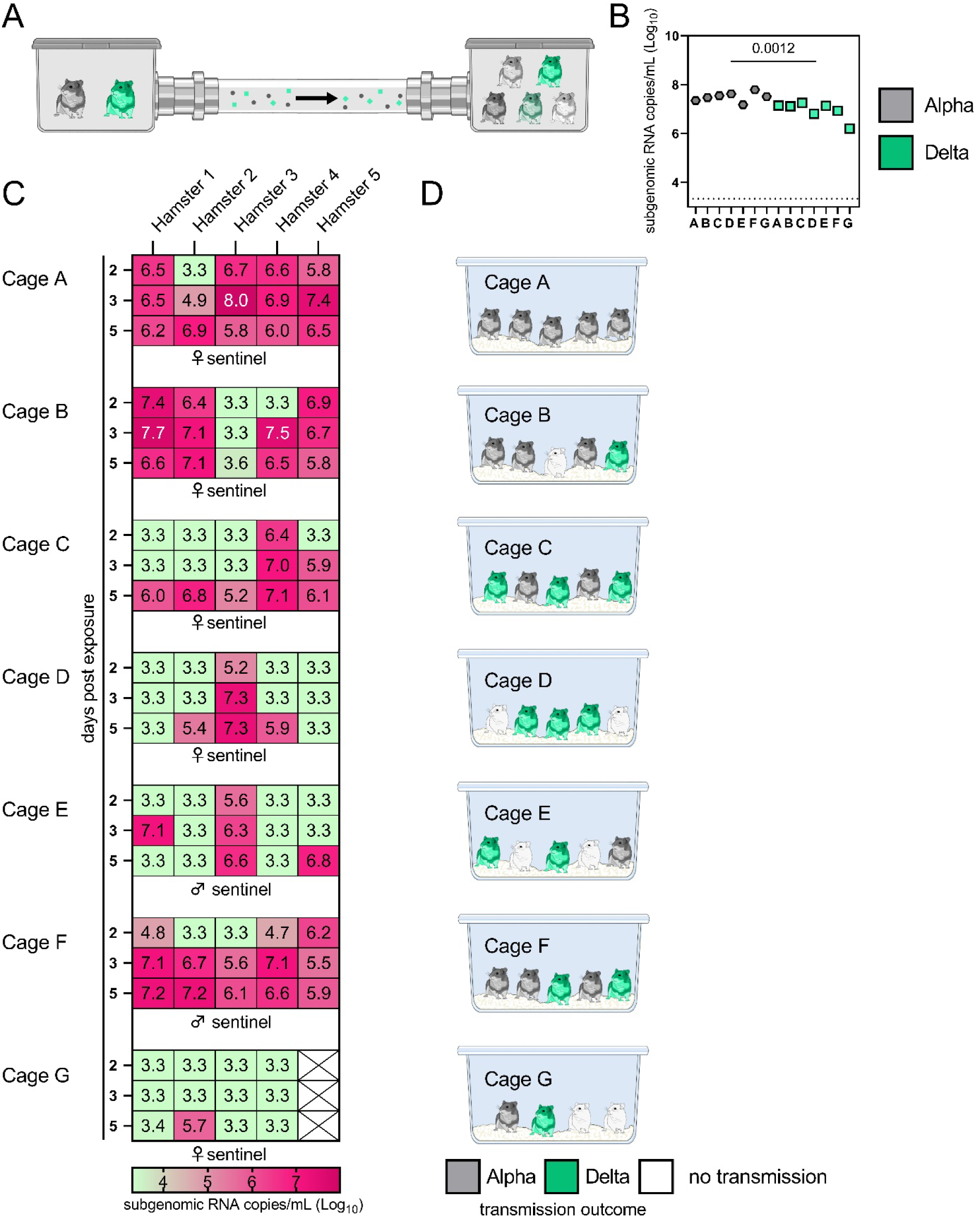
Airborne attack rate of Alpha and Delta SARS-CoV-2 variants. Donor animals (N = 7) were inoculated with either the Alpha or Delta variant with 10^3^ TCID_50_ via the intranasal route and paired together randomly (1:1 ratio) in 7 attack rate scenarios (A-G). To each pair of donors, one day after inoculation, 4-5 sentinels were exposed for a duration of 4 h (i.e., h 24-28 post inoculation) in an aerosol transmission set-up at 200 cm distance. **A.** Schematic figure of the transmission set-up. **B**. Day 1 sgRNA detected in oral swabs taken from each donor after exposure ended. Individuals are depicted. Wilcoxon test, N = 7. Grey = Alpha, teal = Delta inoculated donors**. C.** Respiratory shedding measured by viral load in oropharyngeal swabs; measured by sgRNA on day 2, 3, and 5 for each sentinel. Animals are grouped by scenario. Colors refer to legend below. 3.3 = limit of detection of RNA (<10 copies/rxn). **D.** Schematic representation of majority variant for each sentinel as assessed by percentage of Alpha and Delta detected in oropharyngeal swabs taken at day 2 and day 5 post exposure by deep sequencing. Grey = Alpha, teal = Delta, white = no transmission.

sgRNA in oral swabs taken on 1 DPI varied between animals (**Figure 4B**). Sentinels were either exposed first for 2 h to one variant and then for 2 h to the second (**Figure 4C**, first 4 iterations), or to both variants at the same time for 4 h (last three iterations). Transmission was confirmed by sgRNA in oral swabs collected from all sentinels at 2, 3, and 5 DPE. On 2 DPE, N = 13/34 sentinels were positive for sgRNA in oral swabs, N = 19/34 on 3 DPE and N = 27/34 on 5 DPE. Swabs from 3 DPE and 5 DPE were sequenced, and the percentage of reads mapped to Alpha, and Delta were compared (**Figure 4D**).

All animals had only one variant detectable on day 3. In total, 12 sentinels were infected with Alpha and 7 with Delta by 3 DPE. At 5 DPE, slightly more sentinels shed Alpha (cartoon hamster representation in Figure 4D depicts majority variant for each individual across both sampling days; **Table S 1** lists sequencing results). Interestingly, we observed one superspreading event in iteration A, in which one donor animal transmitted Alpha to all sentinels. For all other iterations, either both donors managed to transmit to at least one sentinel, or not all sentinels were infected. For the iterations with simultaneous exposure, attack rates were similar and statistically indistinguishable: Alpha = 50 %, Delta = 42.8 %. In one simultaneous exposure (iteration F), three sentinels had both Delta and Alpha detectable at 5 DPE. In two, Delta was dominant, and in one Alpha, always with the other variant in the clear minority (<15%). We did not observe any other such coinfections (defined as a PCR positive animal with both Alpha and Delta at 5 % frequency or higher by NGS). This led us to ask whether there was virus interference in sequential exposures -that is, whether established infection with one variant could reduce the probability of successful infection given a later exposure.

To assess this, we used our within-host dynamics model to calculate the estimated infection probabilities for Alpha and Delta for each sentinel in each iteration, assuming each sentinel is exposed independently, but accounting for the different exposure durations, donor sexes, and donor viral load (as measured by oral swabs). From those probabilities, we then calculated posterior probability distributions for the number of co-infections predicted to occur in each iteration if Alpha and Delta infections occurred independently and did not interfere with each other (**Appendix Mathematical Model Figures M2-M4**). We found that our observed coinfections were consistent with this null model; our data do not provide clear evidence of virus interference during sequential exposure, though they also do not rule out such an effect. No difference in virus replication or disease severity was observed between the sentinels infected with Alpha or Delta (**Figure S 6**)

### Limited sustainability of heterologous VOC populations through multiple rounds of airborne transmission

To assess the transmission efficiency in direct competition between the Alpha and Delta VOCs, we conducted an airborne transmission experiment over three subsequent rounds of exposure (**Figure 5A**). Donor animals (N = 8) were inoculated IN with 5 x 10^2^ TCID_50_ of Alpha and 5 x 10^2^ TCID_50_ Delta (1:1 mixture) and eight sentinels were exposed (Sentinels 1, 1:1 ratio) on 1 DPI for 24 h (first chain link, exposure window: 24-48 h post inoculation of the donors) (**Figure 5**). Two days after the start of this exposure, the eight sentinels were placed into the donor side of a new cage and eight new sentinels (Sentinels 2) were exposed for 24 h (second chain link, exposure window 48-72 h post exposure start of the Sentinels 1). Again, 2 days after exposure start, this sequence was repeated for Sentinels 3 (third chain link, exposure window 48-72 h post exposure start of the Sentinels 2). All animals were individually housed between exposures, and after exposure as well for the sentinels. We assessed viral presence in oropharyngeal swabs taken from all animals at 2 and 5 DPI/DPE. While all Sentinels 1 demonstrated active shedding at 2 and 5 DPE, in the Sentinels 2 group no viral RNA was detected in 2/8 animals and no infectious virus in 4/8 by 5 DPE. In the Sentinels 3 group, sgRNA and infectious virus were only detected robustly in one animal on 5 DPE. In contrast to donor animals, all infected sentinels exhibited higher shedding on day 5 compared to day 2 (2 DPI / 5 DPI Donors: median gRNA = 7.8 / 6.9 copies/mL (Log^10^), median sgRNA = 7.2 / 6.4 copies/mL (Log_10_)), median infectious virus titer = 2.3 / 0.5 TCID_50_/mL (Log_10_); Sentinels 1 (median gRNA = 7.2 / 7.4 copies/mL (Log_10_), median sgRNA = 6.4 / 6.9 copies/mL (Log_10_), median infectious virus titer = 2.9 / 2.6 TCID_50_/mL (Log_10_); Sentinels 2 = median gRNA = 3.7 / 5.4 copies/mL (Log_10_), median sgRNA = 1.8 / 3.0 copies/mL (Log_10_), median infectious virus titer = 0.5 / 1.6 TCID_50_/mL (Log_10_)) (**Figure 5B**). Taken together, this evidence suggests that the infectious shedding profile shifts later and decreases in magnitude with successive generations of transmission. This could be explained by lower exposure doses causing lower and slower infections in the recipients.

**Figure 5.**
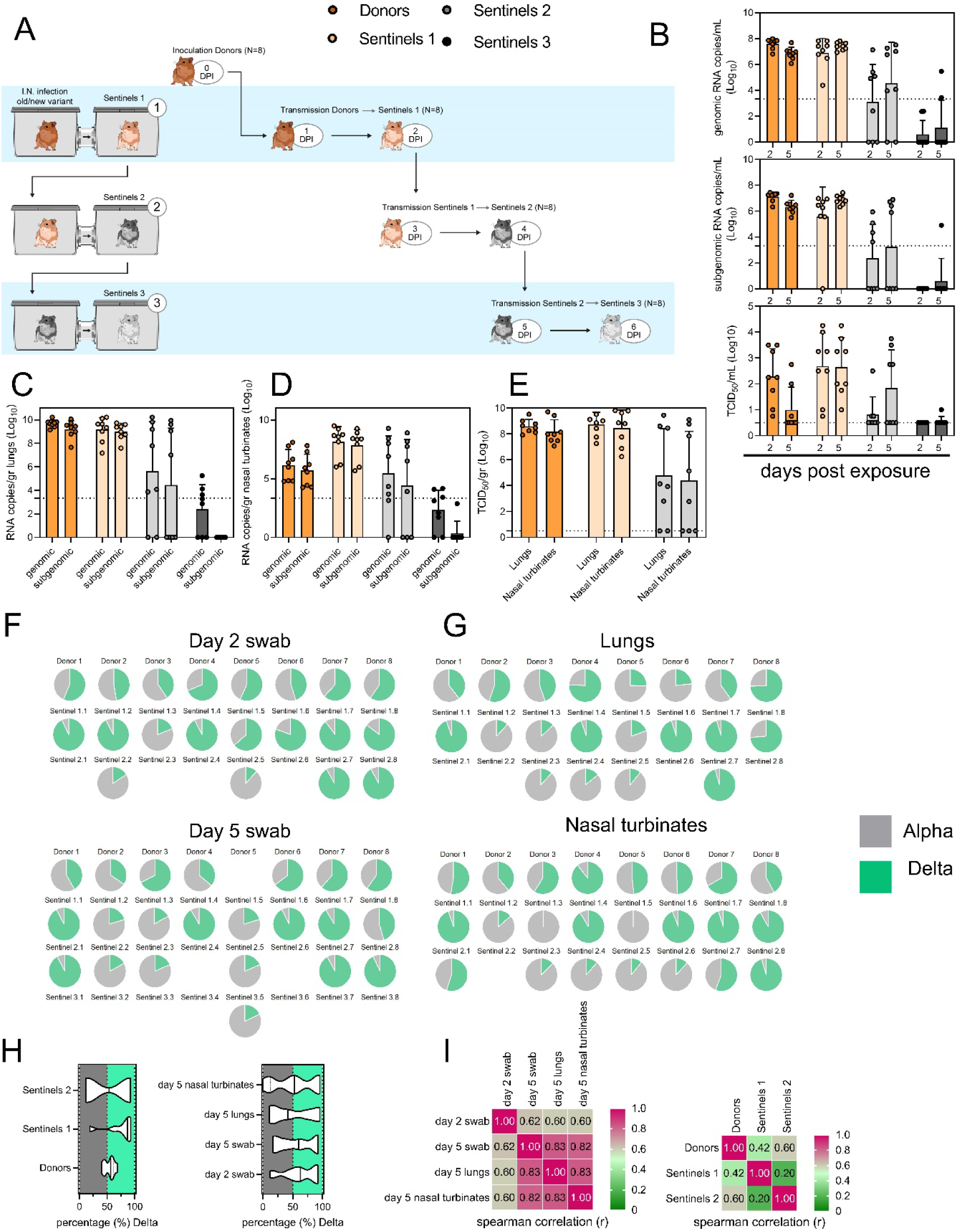
Airborne competitiveness of Alpha and Delta SARS-CoV-2 variants. **A**. Schematic. Donor animals (N = 8) were inoculated with Alpha and Delta variant with 5 x 10^2^ TCID_50_, respectively, via the intranasal route (1:1 ratio), and three groups of sentinels (Sentinels 1, 2, and 3) were exposed subsequently at a 16.5 cm distance. Animals were exposed at a 1:1 ratio; exposure occurred on day 1 (Donors ◊ Sentinels 1) and day 2 (Sentinels ◊ Sentinels). **B.** Respiratory shedding measured by viral load in oropharyngeal swabs; measured by gRNA, sgRNA, and infectious titers on days 2 and day 5 post exposure. Bar-chart depicting median, 96% CI and individuals, N = 8, ordinary two-way ANOVA followed by Šídák’s multiple comparisons test. **C/D/E.** Corresponding gRNA, sgRNA, and infectious virus in lungs and nasal turbinates sampled five days post exposure. Bar-chart depicting median, 96% CI and individuals, N = 8, ordinary two-way ANOVA, followed by Šídák’s multiple comparisons test. Dark orange = Donors, light orange = Sentinels 1, grey = Sentinels 2, dark grey = Sentinels 3, p-values indicated where significant. Dotted line = limit of detection. **F.** Percentage of Alpha and Delta detected in oropharyngeal swabs taken at days 2 and day 5 post exposure for each individual donor and sentinel, determined by deep sequencing. Pie-charts depict individual animals. Grey = Alpha, teal = Delta. **G.** Lung and nasal turbinate samples collected on day 5 post inoculation/exposure. **H**. Summary of data of variant composition, violin plots depicting median and quantiles for each chain link (left) and for each set of samples collected (right). Shading indicates majority of variant (grey = Alpha, teal = Delta). **I.** Correlation plot depicting spearman r for each chain link (right, day 2 swab) and for each set of samples collected across all animals (left). Colors refer to legend on right. Abbreviations: TCID, Tissue Culture Infectious Dose.

We then proceeded to compare the viral loads in the lungs and nasal turbinates at 5 DPE. Viral gRNA was detected in the lungs (**Figure 5C**) and nasal turbinates (**Figure 5D**) of all Donors (lungs: median gRNA = 9.7 copies/gr tissue (Log_10_), nasal turbinates: median gRNA = 6.2 copies/gr tissue (Log_10_)). Interestingly, while the gRNA amount was similar in lungs between Donors and Sentinels 1 (lungs: median gRNA = 9.5 copies/gr tissue (Log_10_)), it was increased in nasal turbinates for the Sentinel 1 group (nasal turbinates: median gRNA = 8.6 copies/gr tissue (Log_10_)). Similarly, sgRNA was increased in Sentinels 1 as compared to Donors in nasal turbinates, but not lungs (Donors = lungs: median sgRNA = 9.4 copies/gr tissue (Log_10_), nasal turbinates: median sgRNA = 5.7 copies/gr tissue (Log10); Sentinels 1 = lungs: median sgRNA = 9.2 copies/gr tissue (Log_10_), nasal turbinates: median sgRNA = 8.4 copies/gr tissue (Log_10_)). Viral gRNA above the level of quantification was detectable in 6/8 Sentinels 2 in both lungs and nasal turbinates, yet sgRNA was only detected in 4/8 Sentinels 2 in lungs and 5/8 in nasal turbinates. Even though gRNA was detected in 3/8 Sentinels 3, no animal had detectable sgRNA in either lungs or nasal turbinates, signaling a lack of active virus replication. To confirm this, infectious virus was analyzed in both tissues for the Donors, Sentinels 1, and Sentinels 2 groups (**Figure 5E**). In both tissues titers were marginally higher in Sentinels 1 (median TCID_50_ / gr tissue (Log_10_) Donors: lungs = 8.6, nasal turbinates = 8.0; Sentinels 1: lungs = 8.9, nasal turbinates = 8.8). Infectious virus was present in 6/8 Sentinels 2 in lungs and 5/8 in nasal turbinates. Hence, even though the exposure interval for the second and third chain links were started 48 h after the start of their own exposure, not all Sentinels 2 became infected, and only one Sentinel 3 animal became infected and demonstrated shedding. We conducted a separate experiment to assess viral loads in the respiratory tract after SARS-CoV-2 airborne transmission at 2 DPI/DPE. While infectious virus was present in oral swabs from all sentinels, virus in lungs and nasal turbinates was not present in all animals (**Figure S 7**).

To determine the competitiveness of the variants, we analyzed the relative composition of the two viruses using next generation sequencing (**Figure 5F,G**). Neither variant significantly outcompeted the other. We first compared the percentage of Delta in oral swabs taken on 2 DPI/DPE, the day of exposure of the next chain link. In Donors, no variant was more prevalent across animals or clearly outcompeted the other within one host (median = 56.5% Delta, range = 40.3-69%). After the first transmission event, Delta outcompeted Alpha at 2 DPE (median = 87.3% Delta, range = 19-92.7%), while after the second transmission event, half (N = 2/4) the animals shed either > 80% either Alpha or Delta. Notably, and in strong contrast to the dual donor experiments described above, every sentinel animal exhibited a mixed infection at 2 DPE, often with proportions resembling those in the donor.

Next, we looked at the selective pressure within the host. By 5 DPI/DPE, no clear difference was observed in Donors (median = 60% Delta, range = 34.3-67.7%), but in the Sentinels 1 group Alpha overtook Delta in three animals (median = 68.3% Delta, range = 17-92.3%), while the reverse was never seen. In one animal, we observed a balanced infection established between both variants at 5 DPE (Sentinel 1.8). In the Sentinels 2 group, Alpha was the dominant variant in N = 3/8 animals, and Delta dominated in 3/8 (median = 55% Delta, range = 17-92.7%). The one Sentinel 3 animal for which transmission occurred shed nearly exclusively Alpha. This suggests that within one host, Alpha was marginally more successful at outcompeting Delta in the oropharyngeal cavity.

We then assessed virus sequences in lungs and nasal turbinates to understand if the selective pressure is influenced by spatial dynamics. In Donor lungs, the percentage of Alpha was marginally higher on 5 DPI (median = 42.3% Delta, range = 23.3-75.7%). In the Sentinels groups, either Alpha or Delta outcompeted the other variant within each animal, only one animal (Sentinel 1.8) revealing both variants > 15%. In N = 5/8 Sentinels 1, yet only in N = 1/4 Sentinel 2 animals, Delta outcompeted Alpha. Sequencing of virus isolated from nasal turbinates reproduced this pattern. In Donors, neither variant demonstrated a completive advantage (median = 51.2% Delta, range = 38.7-89.3%). In N = 5/8 Sentinels 1, and N = 3/8 Sentinels 2, Delta outcompeted Alpha. Combined a trend, while not significant, was observed for increased replication of Delta after the first transmission event, but not after the second, and in the oropharyngeal cavity (swabs) as opposed to lungs (**Figure 5H**) (Donors compared to Sentinels 1: p = 0.0559; Donors compared to Sentinels 2: p = >0.9999; Kruskal Wallis test, followed by Dunn’s test). Swabs taken at 2 DPI/DPE did significantly predict variant patterns in swabs on 5 DPI/DPE (Spearman’s r = 0.623, p = 0.00436) and virus competition in the lower respiratory tract (Spearman’s r = 0.60, p = 0.00848). Oral swab samples taken on day 5 strongly correlate with both upper (Spearman’s r = 0.816, p = 0.00001) and lower respiratory tract tissue samples (Spearman’s r = 0.832, p = 0.00002) taken on the same day (**Figure 5I**).

## Discussion

In immunologically naïve humans, peak SARS-CoV-2 shedding occurs multiple days after exposure and in some cases multiple days before onset of symptoms [26]. It is not known how this informs the window of transmissibility, which is poorly understood and difficult to study in the absence of controlled exposures. Measuring the quantity of exhaled virus and size distribution of airborne particles can provide additional insight into the window of transmissibility beyond simply measuring infectious virus in upper respiratory tract swabs. In addition, the shedding of virus in large and fine aerosols may be a function of physiological changes after infection. Past studies in hamsters have shown that SARS-CoV-2 transmissibility is limited to the first three days. This coincides with peak shedding and ends before the onset of weight loss, clinical manifestation, and loss of infectious virus shedding in the upper respiratory tract [2, 18, 19]. We set out to determine if SARS-CoV-2 infection affects host-derived determinants of airborne transmission efficiency early after infection, which may explain this restriction.

Human studies have found similar peak viral RNA levels for Alpha and Delta [27, 28] despite their epidemiological differences, including Delta’s higher transmissibility [29], shorter generation interval [30], and greater risk of severe disease [31]. We observe similar kinetics in a controlled experimental setting using the hamster model. We found that swab viral load measurements are a valuable imperfect proxy for the magnitude and timing of airborne shedding. Crucially, there is a period early in infection (around 24 h post-infection in inoculated hamsters) when oral swabs show high infectious virus titers, but air samples show low or undetectable levels of virus. Viral shedding should not be treated as a single quantity that rises and falls synchronously throughout the host; spatial models of infection may be required to identify the best correlates of airborne infectiousness [32]. Attempts to quantify an individual’s airborne infectiousness from swab measurements should thus be interpreted with caution, and these spatiotemporal factors should be considered carefully.

While past studies have used whole body plethysmography to differentiate the impact of VOCs on lung function, these have mostly focused on using mathematically derived parameters such as Penh, to compare significant differences on pathology in late acute infection [23]. Within our experimental setup we observed high variation within and between different hamsters. Observed differences could be contributed to the behavioral state which correlated with sex, highlighting that future studies of this nature may require increased acclimatization of the animals to these experimental procedures. However, we did observed changes in breathing patterns as early as 2 DPI, preceding clinical symptoms, but coinciding with the window of time when infectious virus was detected in the air.

The majority of SARS-CoV-2 exhaled from hamsters was observed within droplet nuclei <5 μm in size [27]. We report a rise in <0.53 μm particles and a drop in particles in the 1-10 μm range after infection. One of the caveats of these measurements in small animals is that detected particles may come from aerosolized fomites, and residual dust generated by movement [33]. In our system, we did not detect any particles originating from dead animals or the environment, but we also saw a noticeable reduction of particles across sizes when movement was minimal, or animals were deeply asleep. Considering the individual variability in the lung function data, we did not observe that this shift in particle production was accompanied by a consistent change in either breathing frequency, tidal volume, or minute volume. It remains to be determined how well airway and particle size distribution dynamics in Syrian hamsters model those in humans. Humans with COVID-19 have been shown to exhale fewer particles than uninfected individuals during normal breathing, but not during coughing [34] and fine aerosols have been found to be the major source of virus-loaded droplets. This suggests that a shorter duration of measurable infectious virus in air, as opposed to the upper respiratory tract, could be partially due to early changes in airway constriction and a reduction in exhaled particles of the optimal size range for transmission. The mechanisms involved in the changing aerodynamic particle profile, and the distribution of viral RNA across particle sizes, require further characterization in the hamster model.

Lastly, we compared the transmission efficiency of the Alpha and Delta variants in this system. We did not find a clear transmission advantage for Delta over Alpha in Syrian hamsters, in either an attack rate simulation or when comparing intra- and inter-host competitiveness over multiple generations of airborne transmission. This contrasts sharply with epidemiological observations in the human population, where Delta rapidly replaced Alpha (and other VOCs). The Syrian hamster model may not completely recapitulate all aspects of SARS-CoV-2 virus kinetics and transmission in humans, particularly as the virus continues to adapt to its human host.

Moreover, at the time of emergence of Delta, a large part of the human population was either previously exposed to and/or vaccinated against SARS-CoV-2; that underlying host immune landscape also affects the relative fitness of variants. Our naïve animal model does not capture the high prevalence of pre-existing immunity preasent in the human population and may therefore be less relevant for studying overall variant fitness in the current epidemiological context. Analyses of the cross-neutralization between Alpha and Delta suggest subtly different antigenic profiles [35], and Delta’s faster kinetics in humans may have also helped it cause more reinfections and “breakthrough” infections [36].

Our two transmission experiments yielded different outcomes. When sentinel hamsters were sequentially exposed, first to Alpha and then to Delta, generally no dual infections—both variants detectable—were observed. In contrast, when we exposed hamsters simultaneously to one donor infected with Alpha and another infected with Delta, we were able to detect mixed-variant virus populations in sentinels in one of the cages (Cage F, Appendix Mathematical Model Figures M2, M3). The fact that we saw both single-lineage and multi-lineage transmission events suggests that virus population bottlenecks at the point of transmission do indeed depend on exposure mode and duration, as well as donor host shedding. Notably, our analysis suggests that the Alpha-Delta co-infections observed in the Cage F sentinels could be due to that being the one cage in which both the Alpha and the Delta donor shed substantially over the course of the exposure (Appendix figures S2, S3). Mixed variant infections were not retained equally, and the relative variant frequencies differed between investigated compartments of the respiratory tract, suggesting roles for randomness or host-and-tissue specific differences in virus fitness.

A combination of host, environmental and virus parameters, many of which vary through time, play a role in virus transmission. These include virus phenotype, shedding in air, individual variability and sex differences, changes in breathing patterns, and droplet size distributions. Alongside recognized social and environmental factors, these host and viral parameters might help explain why the epidemiology of SARS-CoV-2 exhibits classic features of over-dispersed transmission [37]. Namely, SARS-CoV-2 circulates continuously in the human population, but many transmission chains are self-limiting, while rarer superspreading events account for a substantial fraction of the virus’s total transmission. Heterogeneity in the respiratory viral loads is high and some infected humans release tens to thousands of SARS-CoV-2 virions/min [38, 39]. Our findings recapitulate this in an animal model and provide further insights into mechanisms underlying successful transmission events. Quantitative assessment of virus and host parameters responsible for the size, duration and infectivity of exhaled aerosols may be critical to advance our understanding of factors governing the efficiency and heterogeneity of transmission for SARS-CoV-2, and potentially other respiratory viruses. In turn, these insights may lay the foundation for interventions targeting individuals and settings with high risk of superspreading, to achieve efficient control of virus transmission [40].”

## Materials and Methods

### Ethics Statement

All animal experiments were conducted in an AAALAC International-accredited facility and were approved by the Rocky Mountain Laboratories Institutional Care and Use Committee following the guidelines put forth in the Guide for the Care and Use of Laboratory Animals 8th edition, the Animal Welfare Act, United States Department of Agriculture and the United States Public Health Service Policy on the Humane Care and Use of Laboratory Animals. Protocol number 2021-034-E. Work with infectious SARS-CoV-2 virus strains under BSL3 conditions was approved by the Institutional Biosafety Committee (IBC). For the removal of specimens from high containment areas virus inactivation of all samples was performed according to IBC-approved standard operating procedures.

### Cells and viruses

SARS-CoV-2 variant Alpha (B.1.1.7) (hCoV320 19/England/204820464/2020, EPI_ISL_683466) was obtained from Public Health England via BEI Resources. Variant Delta (B.1.617.2/) (hCoV-19/USA/KY-CDC-2-4242084/2021, EPI_ISL_1823618) was obtained from BEI Resources. Virus propagation was performed in VeroE6 cells in DMEM supplemented with 2% fetal bovine serum, 1 mM L-glutamine, 50 U/mL penicillin and 50 μg/mL streptomycin (DMEM2). VeroE6 cells were maintained in DMEM supplemented with 10% fetal bovine serum, 1 mM L-glutamine, 50 U/mL penicillin and 50 μg/ml streptomycin. No mycoplasma and no contaminants were detected. All virus stocks were sequenced; no SNPs compared to the patient sample sequence were detected in the Delta stock. In the Alpha stock we detected: ORF1AB D3725G: 13% ORF1AB L3826F: 18%.

### Pseudotype entry assay

The spike coding sequences for SARS-CoV-2 variant Alpha and Delta were truncated by deleting 19 aa at the C-terminus. The S proteins with the 19 aa deletions of coronaviruses were previously reported to show increased efficiency incorporating into virions of VSV [41, 42]. These sequences were codon optimized for human cells, then appended with a 5′ kozak expression sequence (GCCACC) and 3′ tetra-glycine linker followed by nucleotides encoding a FLAG-tag sequence (DYKDDDDK). These spike sequences were synthesized and cloned into pcDNA3.1^+^(GenScript). Human and hamster ACE2 (Q9BYF1.2 and GQ262794.1, respectively) were synthesized and cloned into pcDNA3.1^+^ (GenScript). All DNA constructs were verified by Sanger sequencing (ACGT). BHK cells were seeded in black 96-well plates and transfected the next day with 100 ng plasmid DNA encoding human or hamster ACE2, using polyethylenimine (Polysciences). All downstream experiments were performed 24 h post-transfection. Pseudotype production was carried out as described previously [43]. Briefly, plates pre-coated with poly-L-lysine (Sigma–Aldrich) were seeded with 293T cells and transfected the following day with 1,200 ng of empty plasmid and 400 ng of plasmid encoding coronavirus spike or no-spike plasmid control (green fluorescent protein (GFP)). After 24 h, transfected cells were infected with VSVΔG seed particles pseudotyped with VSV-G as previously described [43, 44]. After one h of incubating with intermittent shaking at 37 °C, cells were washed four times and incubated in 2 mL DMEM supplemented with 2% FBS, penicillin/streptomycin, and L-glutamine for 48 h. Supernatants were collected, centrifuged at 500x*g* for 5 min, aliquoted, and stored at −80 °C. BHK cells previously transfected with ACE2 plasmids of interest were inoculated with equivalent volumes of pseudotype stocks. Plates were then centrifuged at 1200x*g* at 4 °C for 1 h and incubated overnight at 37 °C. Approximately 18–20 h post-infection, Bright-Glo luciferase reagent (Promega) was added to each well, 1:1, and luciferase was measured. Relative entry was calculated by normalizing the relative light unit for spike pseudotypes to the plate relative light unit average for the no-spike control. Each figure shows the data for two technical replicates.

### Structural interaction analysis

The locations of the described spike mutations in the Alpha and Delta VOCs were highlighted on the SARS-CoV-2 spike structure (PDB 6ZGE, [45]). To visualize the molecular interactions at the RBD-ACE2 binding interface, the crystal structure of the Alpha variant RBD and human ACE2 complex (PDB 7EKF [46]) was utilized. All figures were generated using The PyMOL Molecular Graphics System (https://www.schrodinger.com/pymol).

### Aerosol caging

Aerosol cages as described by Port *et al.* [2] were used for transmission experiments and air sampling as indicated. The aerosol transmission system consisted of plastic hamster boxes (Lab Products) connected by a plastic tube. The boxes were modified to accept a 7.62 cm (3’) plastic sanitary fitting (McMaster-Carr), which enabled the length between the boxes to be changed. Airflow was generated with a vacuum pump (Vacuubrand) attached to the box housing the naïve animals and was controlled with a float-type meter/valve (McMaster-Carr).

### Hamster infection with Alpha and Delta

Four-to-six-week-old female and male Syrian hamsters (ENVIGO) were inoculated (10 animals per virus) intranasally (IN) with either SARS-CoV-2 variant Alpha (B.1.1.7) (hCoV320 19/England/204820464/2020, EPI_ISL_683466), variant Delta (B.1.617.2/) (hCoV-19/USA/KY-CDC-2-4242084/2021, EPI_ISL_1823618)., or no virus (anesthesia controls). IN inoculation was performed with 40 µL sterile DMEM containing 10^3^ TCID_50_ SARS-CoV-2 or simply sterile DMEM. At five days post inoculation (DPI), five hamsters from each group were euthanized and tissues were collected. The remaining 5 animals were euthanized at 14 DPI for disease course assessment and shedding analysis. For the control group, no day 5 necropsy was performed. Hamsters were weighed daily, and oropharyngeal swabs were taken on day 1, 2, 3, 4, 5, and 7. Swabs were collected in 1 mL DMEM with 200 U/mL penicillin and 200 µg/mL streptomycin. For the control group, mock swabs were performed to ensure animals underwent the same anesthesia protocols as infection groups. On day −1, 0, 1, 2, 3, 4, 5, 6, 7, and 14 whole body plethysmography was performed. Profiles of particles produced by hamsters were collected on day 0, 1, 3, and 5. Cage air was sampled on day 0, 1, 2, 3, 4, and 5. Hamsters were observed daily for clinical signs of disease. Necropsies and tissue sampling were performed according to IBC-approved protocols.

### Air sampling of hamster cages

During the first 5 days, hamsters were housed in modified aerosol cages (only one hamster box) hooked up to an air pump. Air flow was generated at 30 cage changes/h. Between the cage and the pump a 47 mm gelatin air filter was installed. Filters were exchanged in 24-h intervals. The filters were dissolved in 5 mL of DMEM containing 10% FBS and presence of virus was determined by qRT PCR and plaque assay.

### Aerodynamic particle sizing of exhaled droplets

Two strategies were used to measure the aerodynamic diameter of droplets exhaled by hamsters. SARS-CoV-2 inoculated hamsters or uninfected control animals were placed into a 1.25 L isoflurane chamber. This allowed free movement of the animal in the chamber. The chamber was hooked up with one port to a HEPA filter. The second port was hooked up to a Model 3321 aerodynamic particle sizer spectrometer (TSI). Both chamber and particle sizer were placed into a BSC class II cabinet. Animals remained in the chamber for 5 x 1 min readings. For each set of readings, there were 52 different particle sizes. For each hamster and timepoint, the total number of particles was calculated and the percent of particles in a particular diameter range was derived using this total. RStudio 2021.09.1 Build 372 Ghost Orchid Release, R version 4.1.2 (2021-11-01), Tidyverse R package version 1.3.1 (2021-04-15), and Emmeans R package version 1.7.2 (2022-01-04) were used for the aerodynamic particle size analysis.

To differentiate between particle profiles produced by an awake and moving animal and those produced by a sleeping animal with limited movement, uninfected age-matched hamsters (3 males and 2 female) were acclimatized to being inside a 38.1 mm inside diameter tube hooked up to a particle sizer (**Figure S 5A,B**). Both tube and particle sizer were placed into a BSC class II cabinet. To acclimate the animals to the tube, sunflower seeds were provided to encourage investigation and free entry and exit from the tube. After animals became used to being in the tube, ends were capped as depicted and 5 x 5 min readings were taken. The particle size was measured using a Model 3321 aerodynamic particle sizer spectrometer (TSI). Particle size profiles were analyzed using TSI software. As a control, particles originating from empty enclosures and euthanized animals were recorded and found to be absent

### Aerosol transmission attack rate experiment

Four-to-six-week-old female and male Syrian hamsters (ENVIGO) were used. In this experiment naïve hamsters (sentinels) were exposed to donors infected with either Alpha or Delta in the same aerosol transmission set-up to evaluate the attack rates of both variants. Donor hamsters were infected intranasally as described above with 10^3^ TCID_50_ SARS-CoV-2 (Alpha or Delta, N = 7, respectively) and housed individually. After 24 h, donor animals were placed into the donor cage. 4 or 5 sentinels were placed into the sentinel cage (N = 34, 7 iterations), which was connected to the donor cage by a 2 m tube and exposed for 4 h. Air flow was generated between the cages from the donor to the sentinel cage at 30 cage changes/h. One donor inoculated with Alpha, and one donor inoculated with Delta were randomly chosen for each scenario. Both donors were either placed together into the donor cage, or, alternatively, first one donor was placed into the cage for 2 h, then the other for 2 h. To ensure no cross-contamination, the donor cages and the sentinel cages were never opened at the same time, sentinel hamsters were not exposed to the same handling equipment as donors, and equipment was disinfected with either 70% ETOH or 5% Microchem after each sentinel. Regular bedding was replaced by alpha-dri bedding to avoid the generation of dust particles. Oropharyngeal swabs were taken for donors after completion of the exposure and for sentinels on days 2, 3, and 5 after exposure. Swabs were collected in 1 mL DMEM with 200 U/mL penicillin and 200 µg/mL streptomycin. Donors were euthanized after exposure ended, and sentinels were euthanized on day 5 for collection of lungs. All animals were always single housed outside the exposure window.

### Variant competitiveness transmission chain

Four-to six-week-old female and male Syrian hamsters (ENVIGO) were used. Donor hamsters (N = 8) were infected intranasally as described above with 10^3^ TCID_50_ SARS-CoV-2 at a 1:1 ratio of Alpha and Delta (exact titration of the inoculum for both variants = 503 TCID_50_, 80% Delta sequencing reads). After 12 h, donor animals were placed into the donor cage and sentinels (Sentinels 1, N = 8) were placed into the sentinel cage (1:1) at a 16.5 cm distance with an airflow of 30 cage changes/h as described by Port et al. [2]. Hamsters were co-housed for 24 h. The following day, donor animals were re-housed into regular rodent caging. One day later, Sentinels 1 were placed into the donor cage of new transmission set-ups. New sentinels (Sentinels 2, N = 8) were placed into the sentinel cage at a 16.5 cm distance with an airflow of 30 changes/h. Hamsters were co-housed for 24 h. Then, Sentinels 1 were re-housed into regular rodent caging and Sentinels 2 were placed into the donor cage of new transmission set-ups one day later. New sentinels (Sentinels 3, N = 8) were placed into the sentinel cage at a 16.5 cm distance with an airflow of 30 changes/h. Hamsters were co-housed for 24 h. Then both Sentinels 2 and Sentinels 3 were re-housed to regular rodent caging and monitored until 5 DPE. Oropharyngeal swabs were taken for all animals at 2 and 5 DPI/DPE. All animals were euthanized at 5 DPI/DPE for collection of lung tissue and nasal turbinates. To ensure no cross-contamination, the donor cages and the sentinel cages were never opened at the same time, sentinel hamsters were not exposed to the same handling equipment as donors, and the equipment was disinfected with either 70% EtOH or 5% Microchem after each sentinel. Regular bedding was replaced by alpha-dri bedding to avoid the generation of dust particles.

### Within-host kinetics model

We used Bayesian inference to fit a semi-mechanistic model of within-host virus kinetics and shedding to our data from inoculated hamsters. Briefly, the model assumes a period of exponential growth of virus within the host up to a peak viral load, followed by exponential decay. It assumes virus shedding into the air follows similar dynamics, and the time of peak air shedding and peak swab viral load may be offset from each other by an inferred factor. Decay of RNA may be slower than that of infectious virus by an inferred factor, representing the possibility, seen in our data, that some amplified RNA may be residual rather than representative of current infectious virus levels. We also inferred conversion factors (ratios) among the various quantities, i.e., how many oral swab sgRNA copies correspond to an infectious virion at peak viral load. We fit the model to our swab and cage air sample data using Numpyro [47], which implements a No-U-Turn Sampler [48]. For full mathematical details of the model and how it was fit, including prior distribution choices and predictive checks (Appendix Mathematical Model Figure M1), see **Appendix: Within-host dynamics model and Bayesian inference methods**.

### Whole body plethysmography

Whole body plethysmography was performed on SARS-CoV-2 and uninfected Syrian hamsters. Animals were individually acclimated to the plethysmography chamber (Buxco Electronics Ltd., NY, USA) for 20 minutes, followed by a 5-minute measurement period with measurements taken continuously and averaged over two-second intervals. Initial data was found to have an especially high rejection index (Rinx) for breaths, so was reanalyzed using a custom Buxco formula to account for differences between mice and hamsters. This included expanding the acceptable balance range, the percent change in volume between inhalation and exhalation, from 20-180% to 15-360%. Reanalysis using this algorithm resulted in the Rinx across all hamsters from one day before infection to 5 days post-infection decreasing from 62.97% to 48.65%. The reanalyzed data were then used for further analysis. Each hamster’s individual averages one day prior to infection were used as their baselines for data analysis.

Areas under the curve (AUCs) for each parameter were calculated for each individual hamster based on their raw deviation from baseline at each time point. Either positive or negative peaks were assessed based on parameter-specific changes. Principal component analyses (PCAs) to visualize any potential clustering of animals over the course of infection were performed for each day on raw values for each of the parameters to accurately capture the true clustering with the least amount of data manipulation. PCAs and associated visualizations were coded in R using RStudio version 1.4.1717 (RStudio Team, 2021). The readxl package version 1.3.1 was then used to import Excel data into RStudio for analysis (Wickham and Bryan, 2019). Only parameters that encapsulated measures of respiratory function were included (zero-centered, scaled). The factoextra package version 1.0.1 (Kassambara and Mundt, 2020) was used to determine the optimal number of clusters for each PCA via the average silhouette width method and results were visualized using the ggplot2 package (Wickham, 2016). Correlation plots were generated based on raw values for each lung function parameter using the corrplot package version 0.90 (Wei and Simko, 2021). The color palette for correlation plots was determined using RColorBrewer version 1.1-2 (Neuwirth, 2014).

### Viral RNA detection

Swabs from hamsters were collected as described above. 140 µL was utilized for RNA extraction using the QIAamp Viral RNA Kit (Qiagen) using QIAcube HT automated system (Qiagen) according to the manufacturer’s instructions with an elution volume of 150 µL. For tissues, RNA was isolated using the RNeasy Mini kit (Qiagen) according to the manufacturer’s instructions and eluted in 60 µL. Sub-genomic (sg) and genomic (g) viral RNA were detected by qRT-PCR [49]. RNA was tested with TaqMan™ Fast Virus One-Step Master Mix (Applied Biosystems) using QuantStudio 6 or 3 Flex Real-Time PCR System (Applied Biosystems). SARS-CoV-2 standards with known copy numbers were used to construct a standard curve and calculate copy numbers/mL or copy numbers/g. Limit of detection = 10 copies/rxn.

### Viral titration

Viable virus in tissue samples was determined as previously described [15]. In brief, lung tissue samples were weighed, then homogenized in 1 mL of DMEM (2% FBS). Swabs were used undiluted. VeroE6 cells were inoculated with ten-fold serial dilutions of homogenate, incubated for 1 h at 37°C, and the first two dilutions washed twice with 2% DMEM. For swab samples, cells were inoculated with ten-fold serial dilutions and no wash was performed. After 6 days, cells were scored for cytopathic effect. TCID_50_/mL was calculated by the Spearman-Karber method. To determine titers in air samples, a plaque assay was used. VeroE6 cells were inoculated with 200 µL/well (48-well plate) of undiluted samples, no wash was performed. Plates were spun for 1 h at RT at 1000 rpm. 800 µL of CMC (500 mL MEM (Cat#10370, Gibco, must contain NEAA), 5 mL PenStrep, 7.5 g carboxymethylcellulose (CMC, Cat# C4888, Sigma, sterilize in autoclave) overlay medium was added to each well and plates incubated for 6-days at 37°C. Plates were fixed with 10% formalin overnight, then rinsed and stained with 1% crystal violet for 10 min. Plaques were counted.

### Serology

Serum samples were analyzed as previously described [50]. In brief, maxisorp plates (Nunc) were coated with 50 ng spike protein (generated in-house, purified recombinant) per well. Plates were incubated overnight at 4°C. Plates were blocked with casein in phosphate buffered saline (PBS) (ThermoFisher) for 1 h at room temperature (RT). Serum was diluted 2-fold in blocking buffer and samples (duplicate) were incubated for 1 h at RT. Secondary goat anti-hamster IgG Fc (horseradish peroxidase (HRP)-conjugated, Abcam) spike-specific antibodies were used for detection and KPL TMB 2-component peroxidase substrate kit (SeraCare, 5120-0047) was used for visualization. The reaction was stopped with KPL stop solution (Seracare) and plates were read at 450 nm. The threshold for positivity was calculated as the average plus 3 x the standard deviation of negative control hamster sera.

### MesoPlex Assay

The V-PLEX SARS-CoV-2 Panel 13 (IgG) kit from Meso Scale Discovery was used to test binding antibodies against spike protein of SARS-CoV-2 with 10,000-fold diluted serum obtained from hamsters 14 DPI. A standard curve of pooled hamster sera positive for SARS-CoV-2 spike protein was serially diluted 4-fold. The secondary antibody was prepared by conjugating a goat anti-hamster IgG cross-adsorbed secondary antibody (ThermoFisher) using the MSD GOLD SULFO-TAG NHS-Ester Conjugation Pack (MSD). The secondary antibody was diluted 10,000X for use on the assay. The plates were prepped, and samples were run according to the kit’s instruction manual. After plates were read by the MSD instrument, data was analyzed with the MSD Discovery Workbench Application.

### Virus neutralization

Heat-inactivated γ-irradiated sera were two-fold serially diluted in DMEM. 100 TCID_50_ of SARS-CoV-2 variant Alpha (B.1.1.7) (hCoV320 19/England/204820464/2020, EPI_ISL_683466) or variant Delta (B.1.617.2/) (hCoV-19/USA/KY-CDC-2-4242084/2021, EPI_ISL_1823618) was added. After 1 h of incubation at 37°C and 5% CO_2_, the virus:serum mixture was added to VeroE6 cells. CPE was scored after 5 days at 37°C and 5% CO_2_. The virus neutralization titer was expressed as the reciprocal value of the highest dilution of the serum that still inhibited virus replication. The antigenic map was constructed as previously described [51, 52] using the antigenic cartography software from https://acmacs-web.antigenic-cartography.org. In brief, this approach to antigenic mapping uses multidimensional scaling to position antigens (viruses) and sera in a map to represent their antigenic relationships. The maps here relied on the first SARS-CoV-2 infection serology data of Syrian hamsters. The positions of antigens and sera were optimized in the map to minimize the error between the target distances set by the observed pairwise virus-serum combinations. Maps were effectively constructed in only one dimension because sera were only titrated against two viruses and the dimensionality of the map is constrained to the number of test antigens minus one.

### Next-generation sequencing of virus

Total RNA was extracted from oral swabs, lungs, and nasal turbinates using the Qia Amp Viral kit (Qiagen, Germantown, MD), eluted in EB, and viral Ct values were calculated using real-time PCR. Subsequently, 11 µL of extracted RNA was used as a template in the ARTIC nCoV-2019 sequencing protocol V.1 (Protocols.io -https://www.protocols.io/view/ncov-2019-sequencing-protocol-bbmuik6w) to generate 1st-strand cDNA. Five microliters were used as template for Q5 HotStart Pol PCR (Thermo Fisher Sci, Waltham, MA) together with 10 uM stock of a single primer pair from the ARTIC nCoV-2019 v3 Panel (Integrated DNA Technologies, Belgium), specifically 76L_alt3 and 76R_alt0. Following 35 cycles and 55°C annealing temperature, products were AmPure XP cleaned and quantitated with Qubit (Thermo Fisher Sci) fluorometric quantitation per instructions. Following visual assessment of 1 µL on a Tape Station D1000 (Agilent Technologies, Santa Clara, CA), a total of 400 ng of product was taken directly into TruSeq DNA PCR-Free Library Preparation Guide, Revision D (Illumina, San Diego, CA) beginning with the Repair Ends step (q.s. to 60 µL with RSB). Subsequent cleanup consisted of a single 1:1 AmPure XP/reaction ratio and all steps followed the manufacturer’s instructions including the Illumina TruSeq CD (96) Indexes. Final libraries were visualized on a BioAnalyzer HS chip (Agilent Technologies) and quantified using KAPA Library Quant Kit - Illumina Universal qPCR Mix (Kapa Biosystems, Wilmington, MA) on a CFX96 Real-Time System (BioRad, Hercules, CA). Libraries were diluted to 2 nM stock, pooled together in equimolar concentrations, and sequenced on the Illumina MiSeq instrument (Illumina) as paired-end 2 X 250 base pair reads. Because of the limited diversity of a single-amplicon library, 20% PhiX was added to the final sequencing pool to aid in final sequence quality. Raw fastq reads were trimmed of Illumina adapter sequences using cutadapt version 1.1227, then trimmed and filtered for quality using the FASTX-Toolkit (Hannon Lab, CSHL). To process the ARTIC data, a custom pipeline was developed [53]. Fastq read pairs were first compared to a database of ARTIC primer pairs to identify read pairs that had correct, matching primers on each end. Once identified, the ARTIC primer sequence was trimmed off. Read pairs that did not have the correct ARTIC primer pairs were discarded. Remaining read pairs were collapsed into one sequence using AdapterRemoval [54] requiring a minimum 25 base overlap and 300 base minimum length, generating ARTIC amplicon sequences. Identical amplicon sequences were removed, and the unique amplicon sequences were then mapped to the SARS-CoV-2 genome (<u>MN985325.1</u>) using Bowtie2 [55]. Aligned SAM files were converted to BAM format, then sorted and indexed using SAMtools [56]. Variant calling was performed using Genome Analysis Toolkit (GATK, version 4.1.2) HaplotypeCaller with ploidy set to 2 [57]. Single nucleotide polymorphic variants were filtered for QUAL > 200 and quality by depth (QD) > 20 and indels were filtered for QUAL > 500 and QD > 20 using the filter tool in bcftools, v1.9 [56]. Pie charts were generated using ggplot2 (Wickham, 2016) in R 4.2.1 using RStudio version 1.4.1717 (RStudio Team, 2021).

### Histopathology

Necropsies and tissue sampling were performed according to IBC-approved protocols. Tissues were fixed for a minimum of 7 days in 10% neutral buffered formalin with 2 changes. Tissues were placed in cassettes and processed with a Sakura VIP-6 Tissue Tek on a 12-h automated schedule using a graded series of ethanol, xylene, and ParaPlast Extra. Prior to staining, embedded tissues were sectioned at 5 µm and dried overnight at 42°C. Using GenScript U864YFA140-4/CB2093 NP-1 (1:1000), specific anti-CoV immunoreactivity was detected using the Vector Laboratories ImPress VR anti-rabbit IgG polymer (# MP-6401) as secondary antibody. The tissues were then processed using the Discovery Ultra automated processor (Ventana Medical Systems) with a ChromoMap DAB kit Roche Tissue Diagnostics (#760-159).

### Statistical Analysis

Significance tests were performed as indicated where appropriate for the data using GraphPad Prism 9. Unless stated otherwise, statistical significance levels were determined as follows: ns = p > 0.05; * = p ≤ 0.05; ** = p ≤ 0.01; *** = p ≤ 0.001; **** = p ≤ 0.0001. Exact nature of tests is stated where appropriate.

### Data availability statement

Data deposited in Figshare (10.6084/m9.figshare.20493045) and Github (https://github.com/dylanhmorris/host-viral-determinants).

## Acknowledgements

We would like to thank Ryan Stehlik, Seth Cooley and Shanda Sarchette, and the animal care takers for their assistance during the study. The following reagent was obtained through BEI Resources, NIAID, NIH: SARS-CoV-2 variant Alpha (B.1.1.7) (hCoV320 19/England/204820464/2020, EPI_ISL_683466) and variant Delta (B.1.617.2/) (hCoV-19/USA/KY-CDC-2-4242084/2021, EPI_ISL_1823618). We thank Neeltje van Doremalen, Emmie de Wit, Brandi Williamson, Sujatha Rashid, Ranjan Mukul, Kimberly Stemple, Bin Zhou, Natalie Thornburg, Sue Tong, Stacey Ricklefs, Sarah Anzick for gracefully sharing viruses or propagating and sequencing stocks. We would like to thank Amy Tillman for assistance with the aerodynamic particle data analysis.

## Author Contributions

JRP designed the studies.

JRP, CKY, RJF, MGH, TB, KB performed the experiments, and DHM and JLS developed and analyzed the mathematical models.

JRP, VA, JCR, JES, CIS, KB, CM, KB, JLS, TAS analyzed results.

RPG generated figures.

JRP, DHM, JCR, JLS, VJM wrote the manuscript.

All co-authors reviewed the manuscript. This manuscript has been deposited as a preprint with bioRxiv under a CC0 license for government authors.

