## supplemental files for "Host and viral determinants of airborne transmission of SARS-CoV-2 in the Syrian hamster"

**This PDF file includes:**

Figures S1 to S7
Table S1
SI References

**Other supporting materials for this manuscript include the following:**
SI Mathematical Model Methods Appendix

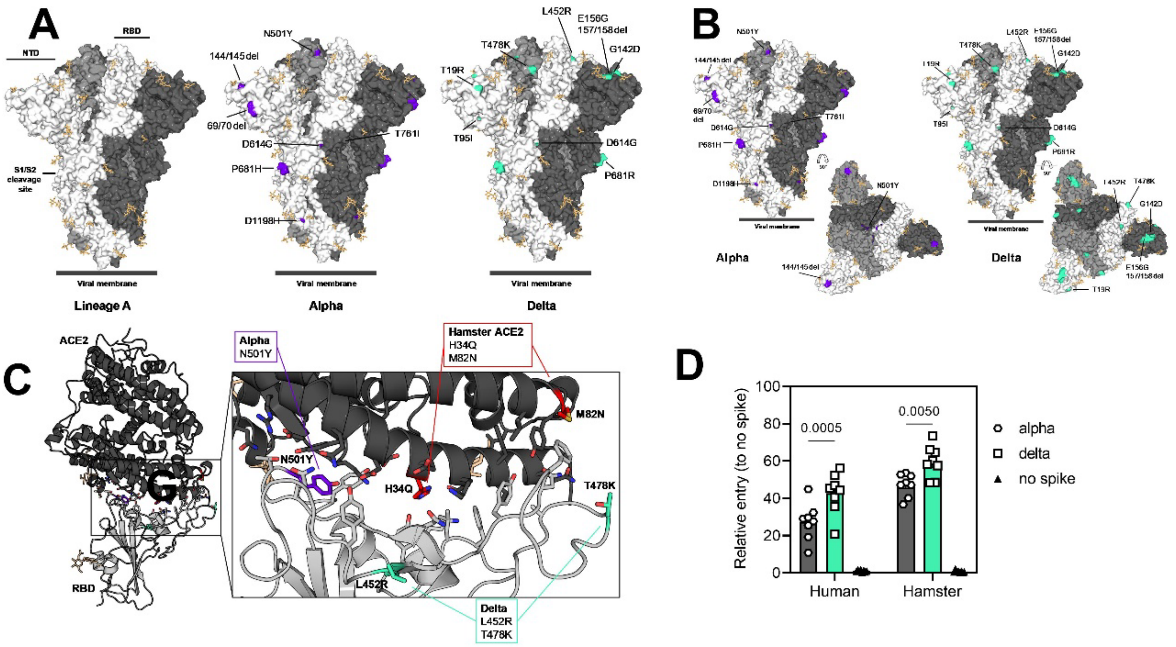


**Figure S1. Alpha and Delta variant spike interaction with hamster ACE2. A/B.** Mutations observed in the SARS-CoV-2 Alpha and Delta VOCs are highlighted on the structure of SARS-CoV-2 spike (PDB 6ZGE, [1]. The spike trimer is depicted by surface representation with each protomer colored a different shade of gray. The residues at the positions of the spike protein mutations observed in the Alpha and Delta SARS-CoV-2 VOCs are colored purple (Alpha) and teal green (Delta) and annotated. N-linked glycans are shown as light, orange-colored sticks. **C.** The structure of the Alpha VOC RBD and human ACE2 complex (PDB 7EKF [2]) is depicted with cartoon representation. ACE2 is colored dark gray and the RBD is colored light gray. N-linked glycans are shown as light, orange-colored sticks. A box reveals a close-up view of the RBD-ACE2 binding interface. Side chains of the residues participating in the interaction, as identified and described by Lan, et al. [3] are shown as sticks. The residues within the RBD that are mutated in the Alpha and Delta VOCs are colored purple (Alpha, N501Y) and teal green (Delta, L452R and T478K). Though they do not participate directly in the ACE2 interface, the sidechains of residues L452 and T478 are also shown. The residues that differ between human and hamster ACE2 within the interface are colored red. **D.** BHK cells expressing either human ACE2 or hamster ACE2 were infected with pseudotyped VSV reporter particles with the spike proteins of Alpha or Delta. Relative entry to no spike control is depicted. Bar-chart depicting median, 95% CI and individuals, N = 8, ordinary two-way ANOVA, followed by Šídák's multiple comparisons test. Abbreviations: RBD, receptor binding domain; ACE2, Angiotensin-converting enzyme 2; VOCs, variants of concern.

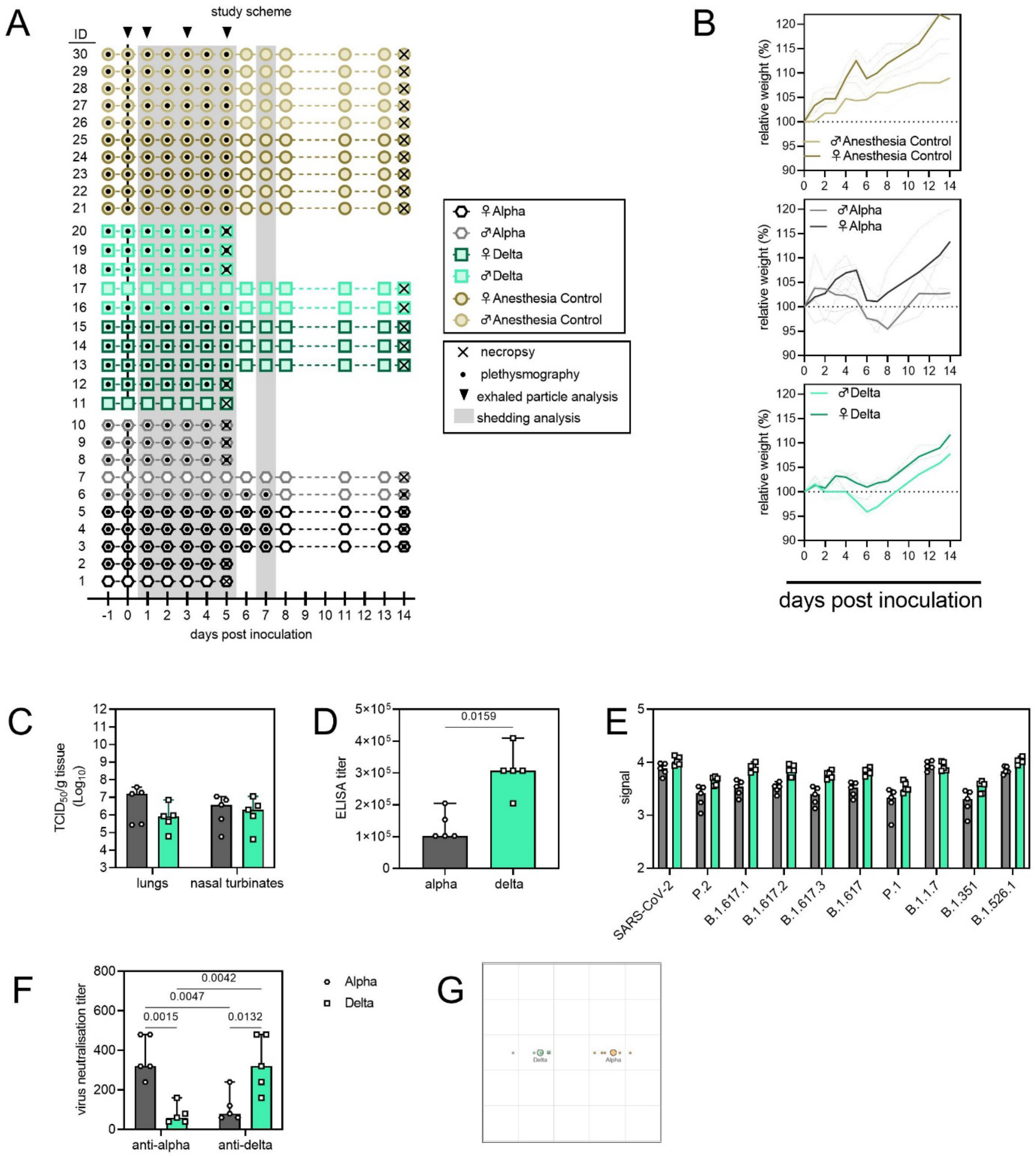


**Figure S2. Schematic overview of inoculation experiment. A**. Four-to-six-week-old female and male Syrian hamsters (ENVIGO) were inoculated (N = 10 per virus, N = 5 per sex) with $10^3$ $TCID_{50}$ intranasally (IN) with either SARS-CoV-2 Alpha or Delta variants, or no virus (anaesthesia controls). At five days post inoculation, five hamsters for each group were euthanized, and tissues were collected. The remaining 5 animals for each route were euthanized at 14 DPI for disease course assessment and shedding analysis. For the control group no day 5 necropsy was performed. Schematic indicates when oropharyngeal swabs were collected, when whole body plethysmography was performed, when air sampling was conducted and when exhaled particle profiles were determined. **B**. Relative weight loss. Graph shows median (thick line) and individuals, colors indicate sex. **C**. Viral load as measured by infectious titers in lungs and nasal turbinates collected at day 5 post inoculation. Bar-chart depicting median, 96% CI and individuals, N = 5, ordinary two-way ANOVA, followed by Šídák's multiple comparisons test. **D**. Binding antibodies against spike protein of SARS-CoV-2 in serum obtained 14 days post inoculation. Bar-chart depicting median, 96% CI and individuals, N = 5, Mann-Whitney test. ELISA was performed once. **E.** Binding antibodies against spike protein of various variants of concern analyzed by MesoPlex. Bar-chart depicting median, 96% CI and individuals, N = 5 ordinary two-way ANOVA, followed by Šídák's multiple comparisons test. Assay was performed once. **F.** Virus neutralization titers against Alpha and Delta, depicted as reciprocal titers. N = 5, ordinary two-way ANOVA, followed by Tukey's multiple comparisons test. Assay was performed once. Grey = Alpha, teal = Delta, beige = anesthesia control. **G.** Antigenic map [4] depicting the cross-reactivity based on neutralization. The spacing between grid lines is 1 unit of antigenic distance, corresponding to a twofold dilution of serum in the neutralization assay. The resulting antigenic distance is depicted between Alpha and Delta. P-values are indicated where significant. Abbreviations: ELISA, Enzyme-linked immune-absorbent Assay.

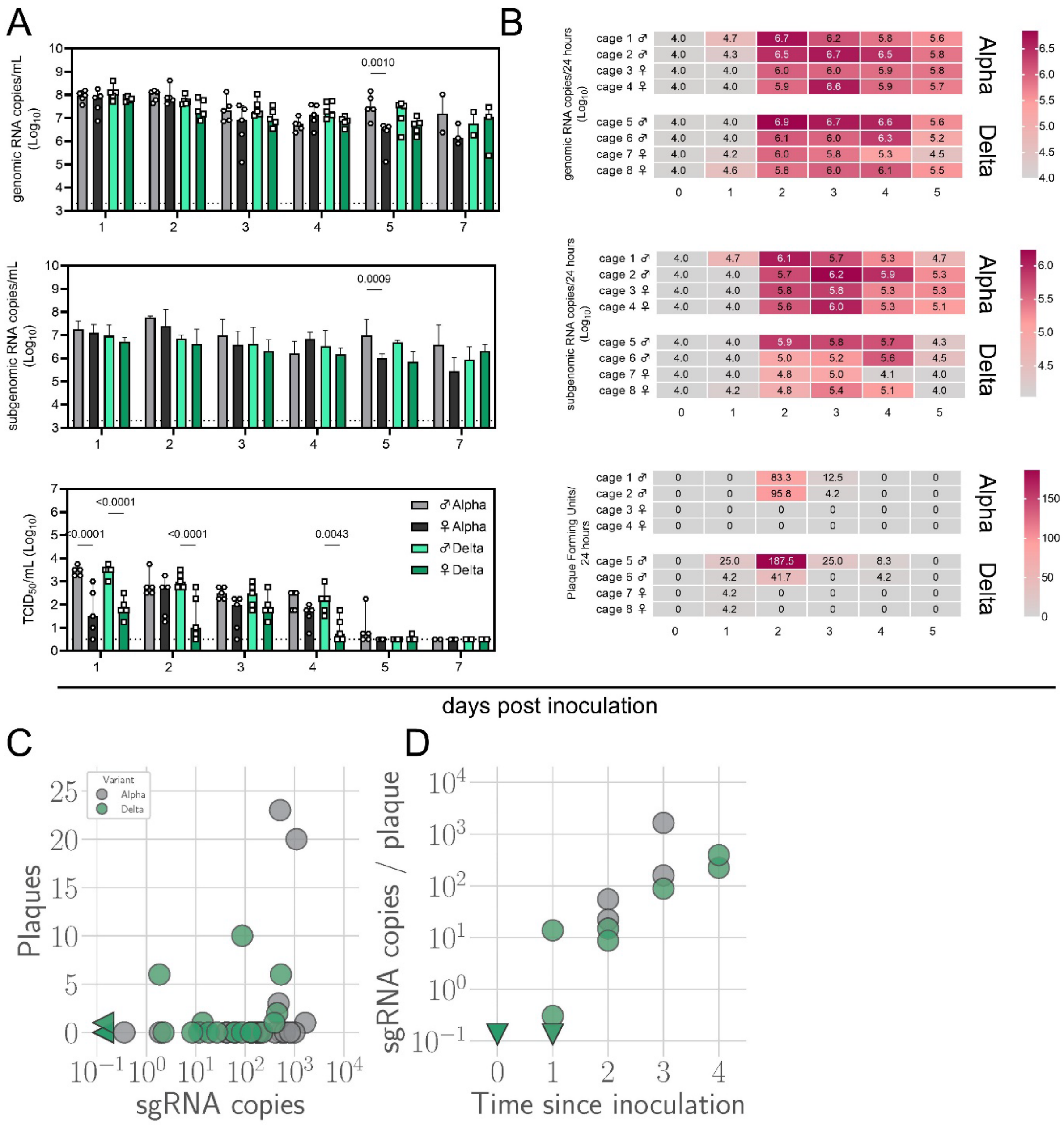


**Figure S3. Window of Alpha and Delta variant shedding profiles.** Syrian hamsters were inoculated with $10^3$ $TCID_{50}$ via the intranasal route with Alpha or Delta. **A**. Viral load as measured by gRNA, sgRNA and infectious titers in oropharyngeal swabs collected at days 1, 2, 3, 4, 5, and 7 post inoculation. Bar-chart depicting median, 96% CI and individuals, N = 5, ordinary two-way ANOVA, followed by Šídák's multiple comparisons test. Dotted line = limit of detection. Grey = Alpha, teal = Delta, dark = female, light = males **B**. Virus isolated from cage air over 24 h intervals, measured as gRNA, sgRNA and plaque forming units on day 0, 1, 2, 3, 4, and 5. The column marked 1 corresponds to samples taken from 0-24 hours post inoculation. Each cage housed 2 or 3 hamsters. Heatmap depicting individual cages across each day, colours referring to legends on the right. RNA: limit of detection = 4.0, Plaque forming units: limit of detection = 0. **C.** sgRNA sampled from air versus infectious virus sampled from air. Point colour indicates variant: Alpha (grey) or Delta (teal). Sampled sgRNA copies value versus sampled plaques. **D.** Number of estimated sgRNA copies per plaque in samples as a function of day sampled and variant. p-values are indicated where significant. Abbreviations: g, genomic; sg, subgenomic; TCID, Tissue Culture Infectious Dose.

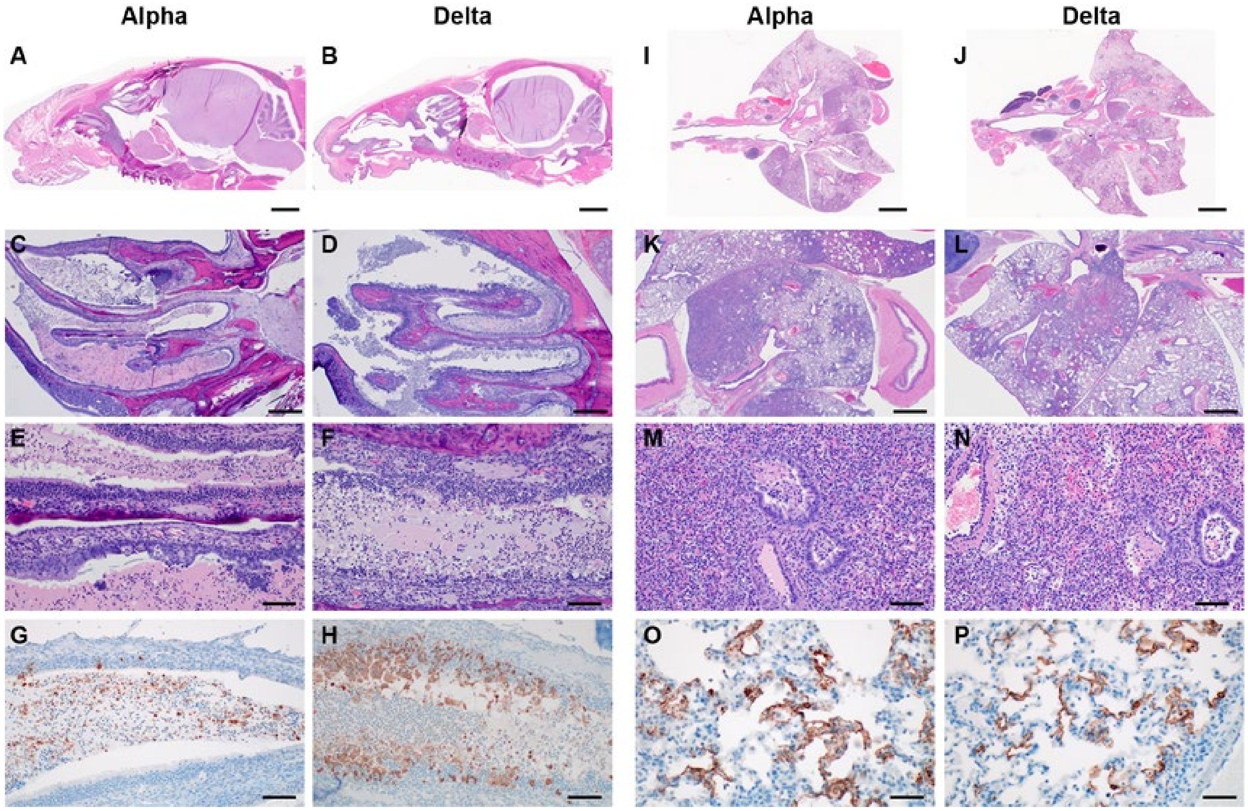

**Figure S4. Respiratory tract pathology after SARS-CoV-2 infection with Alpha and Delta.** Syrian hamsters were inoculated with $10^3$ $TCID_{50}$ via the intranasal route with Alpha or Delta and respiratory pathology was compared on day 5. **A, B.** Skull, brain, nasal turbinates and cavities, Alpha and Delta variant, 1x HE. **C, D**. Nasal cavities contain an exudate and are lined by inflamed, disrupted and necrotic epithelium, 40x HE. **E, F**. Eroded and ulcerated olfactory epithelium, epithelial micro-abscesses, sloughed epithelial and inflammatory exudate occupies nasal cavities and recesses, 200x HE. **G, H**. Examples of immunoreactivity in the exudate of the nasal cavity. There are also rare immunoreactive olfactory epithelial cells in the Alpha variant sample and numerous immunoreactive epithelial cells in the Delta variant sample. 200x, anti-SARS-CoV-2 IHC. **I, J.** Lung, Alpha and Delta variant, 1x HE. **K, L.** Foci of inflammation across multiple lobes, 20x HE. **M, N**. Foci of inflammation contain numerous inflammatory cells as well as hemorrhage, fibrin and edema. Bronchioles contain fibrin, sloughed epithelial and inflammatory cells. Vessels contain sub-intimal leukocytes and vessel walls are occasionally infiltrated by leukocytes, 200x HE. **O, P**. Alveolar immunoreactivity, 400x, anti-SARS-CoV-2 IHC. Mainly, severe inflammation and not yet well-developed interstitial pneumonia were observed. Foci were consolidated but only rarely contained hyperplastic type II pneumocytes and syncytial cells. The large and mid-caliber bronchioles were frequently lined by hyperplastic respiratory epithelium mixed with rare singular necrotic cells and transmigrating leukocytes. Lesions were associated with hemorrhage, fibrin and edema. Vessels also contained sub-endothelial clusters of leukocytes within the muscular layers and surrounding adventitia. In the nasal turbinates SARS-CoV-2 was seen in the exudate of the nasal and rarely in olfactory epithelium, regardless of variant. No sex-specific increases in pathology were observed.

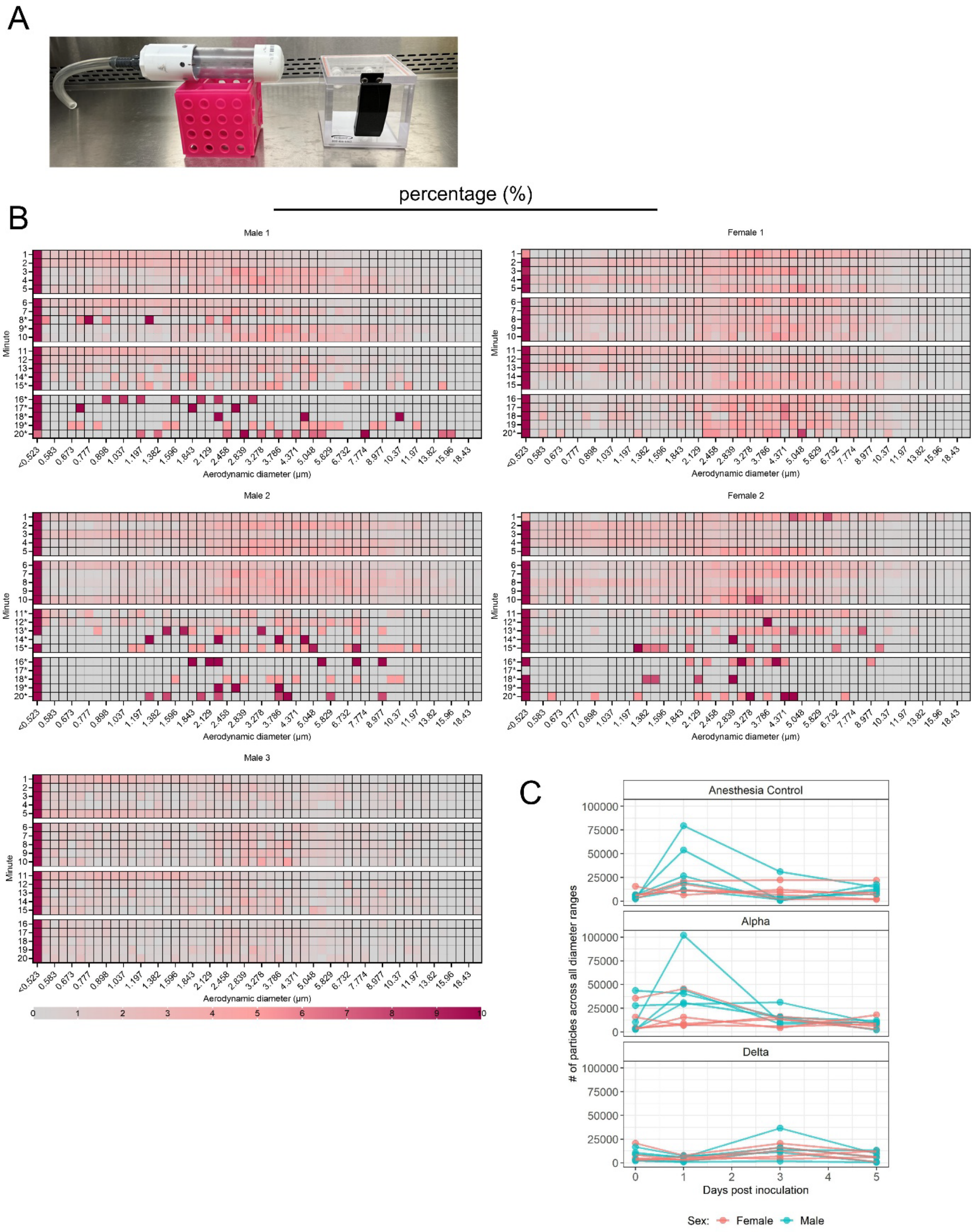


**Figure S5. Exhaled particle profiles of Syrian hamsters A.** Chambers used to house uninfected hamsters (left) and animals during inoculation experiments (inoculation with Alpha, Delta or control inoculum) (right). **B.** Uninfected healthy animals were used to assess particle profiles in relation to behavior patterns. Five Syrian hamsters were acclimatized to a small tube, in which animal movement was limited and air flow was directly passing the face. For each animal, 5 x 5-minute readings were taken. Heatmap shows the percentage of total particles in each diameter size for 3 male and 2 female animals. * marks minutes of low to no activity (sleep). Colors refer to scale below. **C**. Syrian hamsters were inoculated with $10^3$ $TCID_{50}$ via the intranasal route with Alpha (N = 10) or Delta (N = 10). Aerodynamic diameter profile of exhaled particles was analyzed on day 0, 1, 3, and 5. For each animal (N = 10 in each variant group, comprising 5 males and 5 females), line graph of the total number of particles by variant and sex indicated by color (red = female; blue = male).

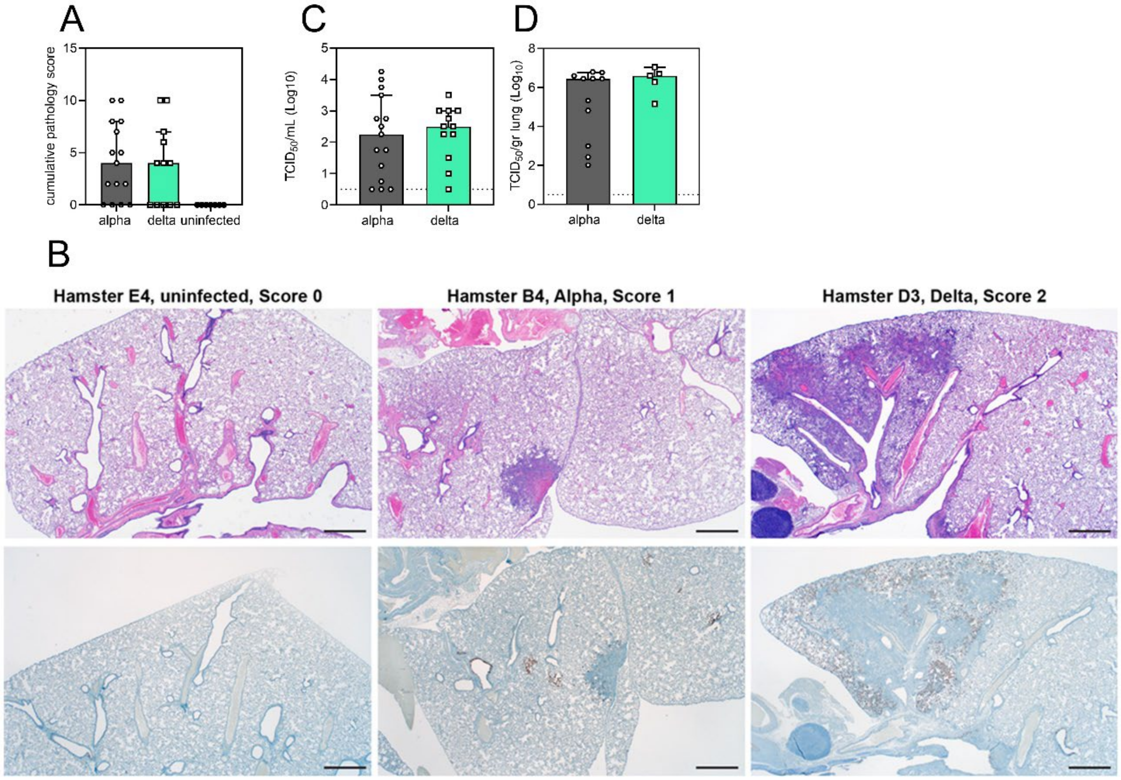


**Figure S6. Airborne competitiveness of Alpha and Delta SARS-CoV-2 variants A.** Cumulative pathology score of sentinels on day 5 post exposure. Bar-chart depicting median, 96% CI, and individuals, Mann-Whitney test. **B**. Lung histologic lesion scores 0, 1, and 2. Score 0, normal lung devoid of immunoreactivity. Score 1, a solitary focus of inflammation (circled) surrounded by normal lung. Bronchiole (*) and alveolar (>) immunoreactivity. Score 2, multiple foci of coalescing inflammation centered on airways. HE (top row) and anti-SARS-CoV-2 IHC (bottom row). **C, D.** Viral load measured via infectious virus titer in swabs and lungs of sentinels on day 5. Bar-chart depicting median, 96% CI, and individuals, Mann-Whitney test. Grey = Alpha, teal = Delta, p-values indicated where significant. Abbreviations: TCID, Tissue Culture Infectious Dose.

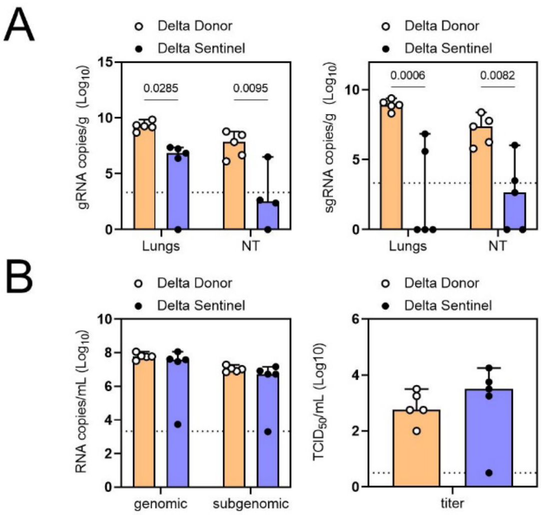


**Figure S7. Early virus shedding in donors and sentinels.** Donor animals were inoculated with Delta variant with $10^3$ $TCID_{50}$ via the intranasal route. Sentinels (1:1 ratio) were exposed subsequently at 16.5 cm distance for 24 h, beginning 24 h after donor exposure. **A.** Organ titers measured by gRNA and sgRNA on day 2 post inoculation/exposure. **B.** Respiratory shedding measured by viral load in oropharyngeal swabs; measured by gRNA, sgRNA and infectious titers on day 2 post inoculation/exposure. Bar-chart depicting median, 96% CI and individuals, N = 5, ordinary two-way ANOVA, followed by Šídák's multiple comparisons test and Wilcoxon test. Orange = donors, purple = sentinels, p-value shown where significant. Abbreviations: g, genomic; sg, subgenomic; TCID, Tissue Culture Infectious Dose.

**Table 1: Airborne attack rate of Alpha and Delta SARS-CoV-2 variants.** Donor animals (N = 7) were inoculated with either the Alpha or Delta variant and paired together randomly in 7 attack rate scenarios (A-G). One day after inoculation, 4-5 sentinels were exposed for a duration of 4 h in an aerosol transmission set-up. Percentage of Alpha and Delta detected in oropharyngeal swabs taken at day 2 and day 5 post exposure by deep sequencing.

**Percentage reads (%) mapped to Delta**

| Cage | Sentinel | Oral swab day | |
|---|---|---|---|
| | | 3 | 5 |
| | A1 | 0 | 0 |
| | A2 | 0 | 0 |
| | A3 | 0 | 0 |
| | A4 | 0 | 0 |
| A | A5 | 0 | 0 |
| | B1 | 0 | 0 |
| | B2 | 0 | 0 |
| | B3 | | |
| | B4 | 0 | 0 |
| B | B5 | 97 | 96.667 |
| | C1 | | 96.333 |
| | C2 | | 0 |
| | C3 | | 96.333 |
| | C4 | 0 | 0 |
| C | C5 | 97.333 | 96.667 |
| | D1 | | |
| | D2 | | 97 |
| | D3 | 96.667 | 97 |
| | D4 | | 97 |
| D | D5 | | |
| | E1 | 97 | |
| | E2 | | |
| | E3 | 97.333 | 97.333 |
| | E4 | | |
| E | E5 | | 0 |
| | F1 | 0 | 0 |
| | F2 | 0 | 0 |
| | F3 | 97 | 90.333 |
| | F4 | 0 | 10.5 |
| F | F5 | 96.667 | 93.667 |
| | G1 | | 0 |
| | G2 | | 97.333 |
| | G3 | | |
| G | G4 | | |
