## appendix for "Host and viral determinants of airborne transmission of SARS-CoV-2 in the Syrian hamster"

### Appendix: Within-host dynamics model and Bayesian inference methods

8.2 Alternative approach to computing viable virions shed during a time window 19

### 1 Introduction

We modeled the observation process explicitly to extract maximal information from the data generated.

#### 1.1 Notation

In the text that follows, we use the following mathematical notation.

##### 1.1.1 Logarithms and exponentials

$\log(x)$ denotes the logarithm base $e$ of $x$ (sometimes called $\ln(x)$). We explicitly refer to the logarithm base 10 of $x$ as $\log_{10}(x)$. $\exp(x)$ denotes $e^x$.

##### 1.1.2 Probability distributions

The symbol $\sim$ denotes that a random variable is distributed according to a given probability distribution. So for example

$$X \sim \text{Normal}(0, 1)$$

indicates that the random variable $X$ is normally distributed with mean 0 and standard deviation 1.

We parameterize normal distributions as:

$$\text{Normal}(\text{mean, standard deviation})$$

We parameterize positive-constrained normal distributions (i.e. with lower limit 0) as:

$$\text{PosNormal}(\text{mode, standard deviation})$$

We parameterize censored normal distributions, in which values outside the censoring range are reported at the lower limit of detection (lld) and upper limit of detection (uld), respectively, as:

$$\text{Censored Normal}(\text{mean, standard deviation, lld, uld})$$

We parameterize Poisson distributions as:

$$\text{Poisson}(\text{mean})$$

##### 1.2 Units

Unless otherwise stated, we express time in units of hours, volume in units of mL, infectious virus in units of plaque forming units collected on our air filter, and sgRNA in units of copy numbers.

#### 2 Dynamics model

We modeled the within-host dynamics of the virus within inoculated hamsters as a process of exponential growth of virus up to a peak, followed by exponential decay of virus down from that peak. The principal quantity of interest is airborne virus shedding over time $V_a(t)$, expressed in units of infectious virions exhaled per unit volume of exhaled air per unit time. We express time in units of hours post-infection.

##### 2.1 Growth and decay of air shedding

We denote the exponential growth rate of the virus within the hamster by $g$ and the exponential decay rate after the peak by $d_{av}$. We denote the time of peak airborne shedding $t_a > 0$ and define $t = 0$ as the time of inoculation.

Our model is therefore:

$$V_a(t) = \begin{cases} V_a(0)\exp\left[gt\right] & t < t_a \\ V_a(0)\exp\left[gt_a - d_{av}(t - t_a)\right] & t \geq t_a \end{cases} \tag{1}$$

##### 2.2 Offset growth and decay of oral shedding

Since we also took measurements of virus shed in oral swabs, we incorporated the dynamics of oral swab shedding $V_o(t)$ into our model. We modeled $V_o(t)$ as offset in time from the dynamics of airborne shedding $V_a(t)$ by some offset factor $\omega > 0$. That is, the time of peak oral shedding $t_o$ is:

$$t_o = \omega t_a \tag{2}$$

Note that $\omega < 1$ implies that swab shedding peaks earlier than airborne shedding, $\omega > 1$ implies that swab shedding peaks later, and $\omega = 1$ implies the peaks coincide in time.

We also allowed for the possibility that oral virus shedding decays at a faster or slower rate $d_{ov}$ than airborne virus shedding, which decays at a rate $d_{av}$. Specifically, we defined the ratio of $d_{ov}$ to $d_{av}$ as $q_o > 0$, so:

$$d_{ov} = q_o d_{av} \tag{3}$$

Then:

$$V_o(t) = \begin{cases} V_o(0)\exp\left[gt\right] & t < t_o \\ V_o(0)\exp\left[gt_o - d_{ov}(t - t_o)\right] & t \geq t_o \end{cases} \tag{4}$$

#### 2.3 Relationship between sgRNA and infectious virus

We also modeled the possibility that measured sgRNA decays slower or faster than measured infectious virus. Slower decay, for instance, could result from persistence of undegraded RNA after all infectious virions have been neutralized or otherwise lost infectivity.

We modeled this possibly different RNA shedding decay rate with an estimated ratio $q_n > 0$ that relates sgRNA shedding decay to infectious virus shedding decay. So the decay rate of airborne sgRNA shedding $d_{an}$ is:

$$d_{an} = q_n d_{av} \tag{5}$$

And similarly the decay rate of oral sgRNA shedding $d_{on}$ is:

$$d_{on} = q_n d_{ov} = q_o q_n d_{av} \tag{6}$$

We modeled the ratio between produced virus $V(t)$ and produced sgRNA copies $N(t)$ with multipliers $o_n$ for oral swabs and $a_n$ for air samples. So before the decay phase begins, $V(t)$ and $N(t)$ are linearly related.

$$\begin{aligned} N_a(t) &= a_n V_a(t) \quad \text{if } t < t_a \\ N_o(t) &= o_n V_o(t) \quad \text{if } t < t_o \end{aligned} \tag{7}$$

The dynamics of airborne sgRNA shedding $N_a(t)$ and oral sgRNA shedding $N_o(t)$ are therefore equivalent to those for infectious virus in equations 1 and 4, respectively, but with $d_{an}$ instead of $d_{av}$, $d_{on}$ instead of $d_{ov}$, and initial values $N_a(0) = a_n V_a(0), N_o(0) = o_n V_o(0)$:

$$N_a(t) = \begin{cases} N_a(0) \exp\left[gt\right] & t < t_a \\ N_a(0) \exp\left[gt_a - d_{an}(t - t_a)\right] & t \geq t_a \end{cases} \tag{8a}$$

$$N_o(t) = \begin{cases} N_o(0) \exp\left[gt\right] & t < t_o \\ N_o(0) \exp\left[gt_o - d_{on}(t - t_o)\right] & t \geq t_o \end{cases} \tag{8b}$$

#### 2.4 Initial shedding value

For inference purposes, rather than set a prior distribution on the initial airborne shedding viral load $V_0 = V_a(0)$, we instead set a prior on the peak airborne viral load $V_{\max} = V_a(t_a)$ and back-calculated $V_a(0)$ (and thus $V_o(0), N_a(0)$, and $N_o(0)$) via:

$$\log(V_0) = \log(V_{\max}) - gt_a \tag{9}$$

#### 2.5 Variant effects

We allowed the two variants of interest, Alpha and Delta, to take on different typical values for all of the virological parameters: $g$, $d_{av}$, $t_a$, $\omega$, $q_o$, $q_n$, $a_n$, $o_n$, and mean peak viral shedding $V_{\max}$. We denote the variant-specific values for variant $i$ by $g_i$, $d_{avi}$, and so on.

#### 2.6 Respiration rates

We also measured animal respiration rates $m(t)$ and included these in our model. The amount of infectious virus an animal deposits into the air per unit time is $m(t)V_a(t)$ and the amount of sgRNA the animal deposits per unit time is $m(t)N_a(t)$.

#### 2.7 Sex effects

Since male hamsters are physically larger and appeared to have different shedding profiles, we wanted to be able to estimate the effect of host sex on key parameters of interest. To do this, we modeled males as possibly offset from females in their typical values of respiration rate $m(t)$, airborne shedding exponential growth rate $g$, airborne shedding exponential decay rate $d_{av}$, peak airborne shedding time $t_a$, and peak airborne shedding rate $V_{\max}$ (this then has downstream consequences for other parameters such as $d_{an}$ or $t_0$ that depend on those core virological parameters).

We modeled sex differences in the virological parameters via offsets to the mean log values for male hamsters. $\Delta_x$ denotes the offset for variable $x$. So for example if females have a mean log respiration rate of $\log[m]$, males have one of $\log[m] + \Delta_m$. We also estimated male offsets $\Delta_g$ for growth rate $g$, $\Delta_d$ for decay rate $d_{av}$, and $\Delta_V$ for peak shedding $V_{\max}$. We did not treat effects as variant-specific, but rather sought to estimate the average sex differences in infection dynamics across the two variants tested.

#### 2.8 Individual heterogeneity in disease course

To account for the fact that individuals have heterogeneous disease courses, we made our model hierarchical, with core virological parameter values for specific individuals distributed about the typical population values. If infected with a variant $i$, animal $j$ has individual values for the virus growth rate $g_{ij}$, the virus decay rate $d_{avij}$, the peak load time $t_{aij}$, and the peak load $V_{\max_{ij}}$.

These values are log-normally distributed about the population values for the given variant and animal sex, with estimated variant-specific standard deviations $\sigma_{gi}$, $\sigma_{di}$, $\sigma_{ti}$ and $\sigma_{Vi}$. We use $s_j$ as an indicator for the sex of hamster $j$ (0 if female, 1 if male). Then:

$$\begin{aligned}
\log[g_{ij}] &\sim \text{Normal}(\log[g_i] + s_j\Delta_g, \sigma_{gi}) \\
\log[d_{avij}] &\sim \text{Normal}(\log[d_{avi}] + s_j\Delta_d, \sigma_{di}) \\
\log[t_{aij}] &\sim \text{Normal}(\log[t_{ai}], \sigma_{ti}) \\
\log[V_{\max_{ij}}] &\sim \text{Normal}(\log[V_{\max_i}] + s_j\Delta_V, \sigma_{Vi})
\end{aligned} \tag{10}$$

We also allowed for individual heterogeneity in respiration rates: animal $j$ has an individual time-averaged respiration rate $m_j$ log-normally distributed about the population value $m$ with an estimated standard deviation $\sigma_m$:

$$\log[m_j] \sim \text{Normal}(\log[m] + s_j\Delta_m, \sigma_m) \tag{11}$$

### 3 Predicting observable quantities

We measured the following quantities:

- Virus subgenomic RNA (sgRNA) in oral swabs, measured via quantitative PCR (qPCR) in units of estimated copy numbers.
- Infectious virus in oral swabs, measured via endpoint titration in units of $\log_{10}$ $\text{TCID}_{50}$ per mL.
- Respiration rates, measured via plethysmography in units of mL air exhaled per unit time.
- Virus sgRNA collected on cage air filters over 24h sampling periods, measured via qPCR in units of estimated copy numbers.
- Infectious virus collected on cage air filters over 24h sampling periods, measured via plaque assay as total plaques formed.
- Infection statuses for each variant for each sentinel hamster.

#### 3.1 Units of virus dynamics

We expressed $V_a(t)$ and $V_o(t)$ in units of total filter-collectible plaque forming units (PFU) shed per mL $\text{h}^{-1}$ (i.e. units that directly predict the cage air infectious virus measurements).

As discussed in section 2, our model explicitly relates infectious virus dynamics $V_a(t)$ and $V_o(t)$ to sgRNA copy number dynamics $N_a(t)$ and $N_o(t)$. The distinct conversion factors $a_n$ and $o_n$ and decay rates $d_{an}$ and $d_{on}$ for airborne versus oral shedding account for two types of possible differences between airborne and oral samples: biological differences (distinct underlying relationships between infectious virus concentration and sgRNA concentration) and measurement differences (distinct quantities of absolute sgRNA quantity recovered given the same underlying sgRNA concentration).

#### 3.2 Converting predicted swab virus to units of TCID50

To fit our model, we needed to convert our internal representation of predicted oral shedding of virus $V_o(t)$, which has the same "predicted air plaques" units as airborne shed virus $V_a(t)$, into the units in which we measured oral shedding: infectious virus $v_o(t)$ in units of $\log_{10}$ $\text{TCID}_{50}/\text{mL}$). We modeled this conversion via a multiplier $o_v$:

$$v_o(t) = o_v V_o(t) \tag{12}$$

The multiplier $o_v$ subsumes both unit conversion and any actual multiplicative difference in virion numbers between airborne shedding and swabs (which could come from lower peak virion concentrations, sampling volume, et cetera.).

#### 3.3 Predicting air sample plaques

We measured shedding into the air at the cage level; multiple hamsters were housed within a single cage. Moreover, samples were cumulative 24h accumulation on the air filter, rather than a point-sample.

##### 3.3.1 Cumulative shedding over a time period

So to fit our model, we needed to compute the cumulative airborne shedding $A$ over some time period $(t_1, t_2)$:

$$A(t_1, t_2) = \int_{t_1}^{t_2} m(t) V_a(t)\, \mathrm{d}t \tag{13}$$

where $m(t)$ is the animal's respiration rate. Integrating yields:

$$A(t_1, t_2) = \begin{cases} \frac{m}{g}\left(V_a(t_2) - V_a(t_1)\right) & t_2 \leq t_a \\ \frac{m}{d_{av}}\left(V_a(t_1) - V_a(t_2)\right) & t_1 > t_a \\ \frac{m}{g}\Big(V_a(t_a) - V_a(t_1)\Big) + \frac{m}{d_{av}}\left(V_a(t_a) - V_a(t_2)\right) & \text{otherwise} \end{cases} \tag{14}$$

where $m$ is an appropriately-chosen constant to represent the time-varying effect of $m(t)$ on the value of the integral. Note that while ideally we would know how $m(t)$ and $V_a(t)$ change together and compute the integral explicitly, in practice we could only measure $m(t)$ coarsely, and so it was simpler to infer the appropriate value, understanding that it would not necessarily equal a naive temporal average.

##### 3.3.2 Accounting for loss of virion infectivity

Furthermore, since the air shedding measured accumulation on the air filter over a 24 hour period, we had to account for decay of infectious virus between exhalation and quantification. To do this, we assumed that the virus decays exponentially in suspended aerosols and on the filter (as we have previously measured empirically[1,2]) but that minimal virus is lost once the filter is removed for virus quantification.

Suppose the the sampling period begins at a time $t_1$ post-infection and ends at a time $t_2$ when the filter is removed. Each hamster $j$ sheds infectious virus at a rate $mV_{aj}(t)$ per unit time. But if the virus loses infectivity according to an exponential decay process with rate $\lambda$, then only a fraction $e^{-\lambda(t_s - t)}$ of the virions shed at time $t_s > t_1$ and collected on the filter will remain infectious when the filter is collected at $t_2$. For simplicity in mathematical notation, we assume here that, as in our experiments, potentially shedding individual remained present until at least $t_2$, though our code allows for modeling other situations, such as the removal of the shedding individual prior to the removal of the filter.

We denote the cumulative number of virions shed since $t_1 \geq t_0$ that remain viable (infectious) at $t_2 \geq t_1$ by $A_v(t_1, t_2)$ (to distinguish it from the cumulative shedding irrespective of retained infectiousness $A$). Fixing $t_1$, $A_v$ is defined by the following differential equation with respect to $t_2$:

$$\frac{\mathrm{d}A_v}{\mathrm{d}t_2} = mV_a(t_2) - \lambda A_v \tag{15}$$

Since we have an expression for $V_a(t)$ consisting of fixed periods of constant-rate exponential growth or decay, we can treat this as a system of two coupled linear ordinary differential equations, with $\frac{\mathrm{d}V_a}{\mathrm{d}t_2} = gV_a$ or $\frac{\mathrm{d}V_a}{\mathrm{d}t_2} = -d_{av}V_a$, depending on whether $t_2 > t_a$, and solve.

Assume the shedding individual is removed at a time $t_s \geq t_1$, which may or may not be after $t_a$. Then:

$$A_v(t_1, t_2) = \begin{cases} \frac{mV(t_1)}{g+\lambda}\left(e^{g(t_2-t_1)} - e^{-\lambda(t_2-t_1)}\right) & t_1 < t_2 \leq t_a, t_s \\ \frac{mV(t_1)}{-d_{av}+\lambda}\left(e^{-d_{av}(t_2-t_1)} - e^{-\lambda(t_2-t_1)}\right) & t_a \leq t_1 < t_2 \leq t_s \\ \frac{mV(t_a)}{-d_{av}+\lambda}\left(e^{-d_{av}(t_2-t_a)} - e^{-\lambda(t_2-t_a)}\right) + A_v(t_1, t_a)e^{-\lambda(t_2-t_a)} & t_1 < t_a < t_2 \leq t_s \\ A_v(t_1, t_s)e^{-\lambda(t_2-t_s)} & t_1 \leq t_s < t_2 \end{cases} \tag{16}$$

In a previous version of this work, we took an equivalent alternative approach based on integrating over the virions shed at a time $t_1 \leq t \leq t_2$ that would survive until $t_2$; we describe that alternative approach in section 8.2 below for completeness.

#### 3.4 Predicting air sample sgRNA

For air sample sgRNA, we again predicted cumulative accumulation. We did not model environmental degradation of detectable sgRNA, but rather chose to treat it implicitly via the decay rate ratio $q_n$ relating airborne infectious virus shedding to airborne sgRNA shedding. We chose not to model sgRNA environmental degradation more explicitly because environmental half-lives for sgRNA are less well-characterized, but likely longer, than environmental half-lives for infectious virus.

The cumulative predicted number of sgRNA copies collected is:

$$A_n(t_1, t_2) = m\int_{t_1}^{t_2} N_a(t)\,\mathrm{d}t \tag{17}$$

$A_n$ can be computed using the following antiderivative:

$$G(t, a, t_i) = \int \exp[a(t - t_i)]\,\mathrm{d}t = \frac{1}{a}\exp[a(t - t_i)] \tag{18}$$

We obtain:

$$A_n(t_1, t_2) = \begin{cases} mN_a(t_1)\left[G(t_2, g.t_1) - G(t_1, g, t_1)\right] & t_2 < t_a \\ \\ mN_a(t_1)\left[G(t_2, -d, t_1) - G(t_1, -d, t_1)\right] & t_1 > t_a \\ \\ mN_a(t_1)\left[G(t_a, g, t_1) - G(t_1, g, t_1)\right] + & t_1 \leq t_a \leq t_2 \\ mN_a(t_a)\left[G(t_2, -d, t_a) - G(t_a, -d, t_a)\right] & \end{cases} \tag{19}$$

#### 3.5 Predicting sentinel exposures

Using our kinetics model, we were able to estimate the probability each donor in our dual donor experiment had of infecting each sentinel, taking into account donor sex, infecting variant, and timing of exposure. This also enabled us to assess whether the absence of observed co-infections in sequential donor experiments was more suggestive of competitive interference or non-interference among the virus variants (see section 6 for methods and results).

To do this, we assumed that each sentinel's dose from each donor was proportional to the cumulative airborne shedding by the donor over the period of sentinel exposure. Given the short exposure period, we ignored the effect of environmental loss, so the computation was $A_v(t_1, t_2)$ as in equation 43, but with the environmental loss rate set to $\lambda = 0$. We assumed that the total dose received by each sentinel was equal to the cumulative shedding multiplied by an estimated variant-specific constant $c_i$ that subsumes uncertainty about sentinel respiration rate, cage airflow, and per-virion infectivity when inhaled by a hamster (since $A_v$ has units of predicted *cell culture* plaques, and hamster airways may be more or less susceptible to virions of a given variant).

So in our model, each sentinel $j$ receives a dose $h_{ij}$ of variant $i$ that depends on the virus shedding $A_{vij}(t_{1ij}, t_{2ij})$ from the donor animal associated to variant $i$ and sentinel $j$, where $t_{0ij}$ and $t_{1ij}$ are the start and end times of the exposure under the exposure design for variant $i$ and sentinel $j$:

$$h_{ij} = c_i A_{vij}(t_{1ij}, t_{2ij}) \tag{20}$$

We again applied a Poisson single-hit model of infection, so the probability $p_{\text{inf}}(i, j)$ that sentinel $j$ is infected with variant $i$ depends on the cumulative dose $h_{ij}$ as:

$$p_{\text{inf}}(i, j) = 1 - e^{-h_{ij}} \tag{21}$$

### 4 Relating predicted quantities to observed quantities

#### 4.1 Oral swabs

##### 4.1.1 Infectious virus titers

Denote the $k^{\text{th}}$ measured oral swab titer by $y_{vok}$. Suppose it was sampled from individual animal $j$ at time $t$. Then its predicted value is $v_{ok} = o_v V_{oj}(t)$.

We modeled the distribution of observed swab titers given $\log_{10}$TCID$_{50}$ values $v_{ok}$ via a Poisson single-hit process, as we have described previously[2,3] (in particular, see section 2.2 of SI of[3]). Briefly, in a Poisson single-hit model of virus titration, a Poisson-distributed number of virions successfully infect cells in each. The mean $\mu$ of this Poisson depends on the underlying sample virus concentration $v$ (in $\log_{10}$TCID$_{50}$) and degree of dilution $D$ (in $\log_{10}$ fold dilutions).

$$\mu = \log(2)10^{v-D} \tag{22}$$

The factor of $\log(2)$ converts from units of TCID$_{50}$ to units of successful virions.

A well will be positive for infection if at least one virion infects a cell, which occurs with probability:

$$1 - \exp\left(-\log(2)10^{v-D}\right) \tag{23}$$

A complication to the typical single-hit model in this case is that we only had total counts of positive wells rather than exact well identities, dilutions, and positive/negative status. To handle this, we used an approximate method that integrates the likelihood function over the most probable configurations of positive and negative wells that could generate an observed total count. We describe this method in section 8.1.

###### 4.1.2 Subgenomic RNA

If $y_{nok}$ is the $k^{\text{th}}$ measurement of oral swab sgRNA, sampled from animal $j$ at time $t$, its predicted value is $n_{ok} = N_o(t)$. To account for different sampling procedures and qPCR runs for the donor animals used in the dual donor experiments compared to the animals used in the kinetics experiments, for the donor animals we added an estimated offset term $f$ to the log copy number: $\log[n_{ok}] = \log[N_o(t)] + f$.

We modeled the observed $\log_{10}$ oral swab sgRNA copy numbers $y_{nok}$ as distributed about their predicted values $n_{ok}$, with an estimated variant-specific standard deviation $\sigma_{noi}$ (where $i$ is the variant infecting animal $j$) and censoring at the minimum and maximum observable values (which are given by the particular sgRNA standard curve).

$$\log_{10}(y_{noj}) \sim \text{Censored Normal}(\log_{10}(n_{oj}), \sigma_{noi}, n_{\text{min}}, n_{\text{max}}) \tag{24}$$

##### 4.2 Air samples

###### 4.2.1 Plaques

We used equations 45, 46, and 48 to predict the number of plaques $v_{ak}$ observed on each filter.

Note that this implies $V_a(t)$ has units of filter plaques produced per mL exhaled air per unit time (in the absence of environmental decay).

If an observed plaque count $y_{avk}$ comes from an air sample taken between time $t_1$ and time $t_2$ from a cage with $n_h$ hamsters infected with variant $j$, the corresponding predicted plaque count $v_{ak}$ is:

$$v_{ak} = \sum_{u=1}^{n_h} A_{vu}(t_1, t_2) \tag{25}$$

where $A_{vu}(t_1, t_2)$ is $A_v(t_1, t_2)$ for the $u^{\text{th}}$ hamster.

Since *in vitro* cell infection is well-described by a Poisson single-hit process[4] (possibly with binomial thinning), we modeled the observed plaque counts $y_{avk}$ as Poisson-distributed about their predicted values $v_{ak}$.

$$y_{vak} \sim \text{Poisson}(v_{ak}) \tag{26}$$

##### 4.2.2 Subgenomic RNA

Similarly, we used equation 17 to predict the number of sgRNA copies $n_{ak}$ that would be observed when sampling cage $k$ from $t_1$ until $t_2$ as:

$$n_{ak} = \sum_{u=1}^{n_h} A_{nu}(t_1, t_2) \tag{27}$$

where $A_{nu}$ is the cumulative sgRNA shedding function $A_n$ for hamster $u$.

We model the observed $\log_{10}$ air sample sgRNA copy numbers $\log_{10}(y_{nak})$ as normally distributed about their predicted values $\log_{10}(n_{ak})$ with an estimated variant-specific standard deviation $\sigma_{nai}$ and censoring at the minimum and maximum possible $\log_{10}$ estimated copy numbers (which depend on the standard curve).

$$\log_{10}(y_{nak}) \sim \text{Censored Normal}(\log_{10}[n_{ak}], \sigma_{nai}, n_{\min}, n_{\max}) \tag{28}$$

#### 4.3 Respiration rates

We modeled the observed log respiration rates for animal $j$ $\log(y_{mij})$ as distributed about the animal's typical log value $\log(m_j)$ with a estimated standard deviation $\sigma_r$.

$$\log(y_{mij}) \sim \text{Normal}(\log(m_j), \sigma_r) \tag{29}$$

#### 4.4 Sentinel infection status

Our dynamical model generates predicted infection probabilities $p_{\text{inf}}(i, j)$ for sentinel $i$ with variant $j$ (see section 3.5).

The observed infection status for sentinel $j$ with variant $i$, $y_{pij} \in \{0, 1\}$ is therefore Bernoulli distributed with probability $p_{\text{inf}}[i, j]$.

$$y_{pij} \sim \text{Bernoulli}(p_{\text{inf}}[i, j]) \tag{30}$$

### 5 Prior distributions

In general, we sought to set prior distributions for our parameters that were "weakly informative"[5]; that is, that rule out biologically implausible or impossible values while remaining fairly agnostic about possible values of interest. We assessed the robustness of our prior distribution choices via prior predictive checks.

#### 5.1 Respiration rates

We placed a normal prior on the population-wide mean log respiration rate $\log(m)$, with $m$ in units of $\text{mL h}^{-1}$.

$$\log(m) \sim \text{Normal}(\log[4800], 0.25) \tag{31}$$

We placed a positive-constrained normal prior on the individual respiration rate standard deviation $\sigma_{mi}$ (see equation 10).

$$\sigma_{mi} \sim \text{PosNormal}(0, 0.25) \tag{32}$$

#### 5.2 Virological parameters

We placed log-normal priors on the variant-specific time to peak $t_{ai}$ and peak shedding rate $V_{\text{max}_i}$; $i$ indexes the variant. To encode prior information about the variant-specific growth and decay rates $g_i$ and $d_{avi}$ in an interpretable manner, we placed normal priors on the doubling and halving times (in hours) $t_{2i} = \log(2)/g_i$ and $t_{\frac{1}{2}i} = \log(2)/d_{avi}$ and then back-calculated $g_i$ and $d_{avi}$.

$$\begin{aligned} \log[t_{ai}] &\sim \text{Normal}(\log[24], 0.5) \\ \log[V_{\text{max}_i}] &\sim \text{Normal}(\log[1] - \log[24] - \log[4800], 3) \\ \log[t_{2i}] &\sim \text{Normal}(\log[5], 0.5) \\ \log[t_{\frac{1}{2}i}] &\sim \text{Normal}(\log[15], 0.75) \\ \log[t_{\frac{1}{2}i}] &\sim \text{Normal}(\log[15], 0.75) \end{aligned} \tag{33}$$

The prior mode for $\log[V_{\text{max}_i}]$ can be interpreted as corresponding to the amount of shedding that would lead to 1 plaque(s) on the air filter from a 24h sample at the prior mean respiration rate of $\log[4800\text{mL h}^{-1}]$.

We placed normal priors on the male sex effects $\Delta_m$, $\Delta_g$, $\Delta_d$, and $\Delta_V$ that modify the virological parameters.

$$\begin{aligned} \Delta_m &\sim \text{Normal}(0, 0.25) \\ \Delta_g &\sim \text{Normal}(0, 0.25) \\ \Delta_d &\sim \text{Normal}(0, 0.25) \\ \Delta_V &\sim \text{Normal}(0, 0.25) \end{aligned} \tag{34}$$

We placed lognormal priors priors on the swab to air peak timing ratio $\omega$, the swab to air decay rate ratio $q_o$, the sgRNA to infectious virus decay rate ratio $q_n$, the air infectious virus to oral TCID conversion factor $o_v$, the air and swab copy number to infectious virus ratios

$a_n$ and $o_n$. We placed a normal prior on the donor log copy number offset $f$.

$$
\begin{aligned}
\log(\omega) &\sim \text{Normal}(0, 0.25) \\
\log(q_o) &\sim \text{Normal}(0, 1) \\
\log(q_n) &\sim \text{Normal}(\log[1], 1) \\
\log(o_v) &\sim \text{Normal}(0, 10) \\
\log(a_n) &\sim \text{Normal}(0, 10) \\
\log(o_n) &\sim \text{Normal}(0, 10) \\
f &\sim \text{Normal}(0, 1.5)
\end{aligned}
\tag{35}
$$

We placed positive-constrained normal priors on the hierarchical standard deviations that specify degree of individual variation about these population-wide virological parameters.

$$
\begin{aligned}
\sigma_{gi} &\sim \text{PosNormal}(0, 0.2) \\
\sigma_{di} &\sim \text{PosNormal}(0, 0.2) \\
\sigma_{ti} &\sim \text{PosNormal}(0, 0.15) \\
\sigma_{Vi} &\sim \text{PosNormal}(0, 2)
\end{aligned}
\tag{36}
$$

#### 5.3 Sentinel infection process constant

We placed a lognormal prior on the variant-specific sentinel infection process constant $c_i$.

$$
\log[c_i] \sim \text{Normal}(0, 3) \tag{37}
$$

#### 5.4 Observation error standard deviations

We placed positive-constrained normal priors on the observation process standard deviations for respiration rate $\sigma_r$ (equation 29), oral swab sgRNA copies $\sigma_{noi}$ (equation 24), and air sample sgRNA copies $\sigma_{nai}$.

$$
\begin{aligned}
\sigma_r &\sim \text{PosNormal}(0, 0.2) \\
\sigma_{noi} &\sim \text{PosNormal}(0, 0.5) \\
\sigma_{nai} &\sim \text{PosNormal}(0, 0.5)
\end{aligned}
\tag{38}
$$

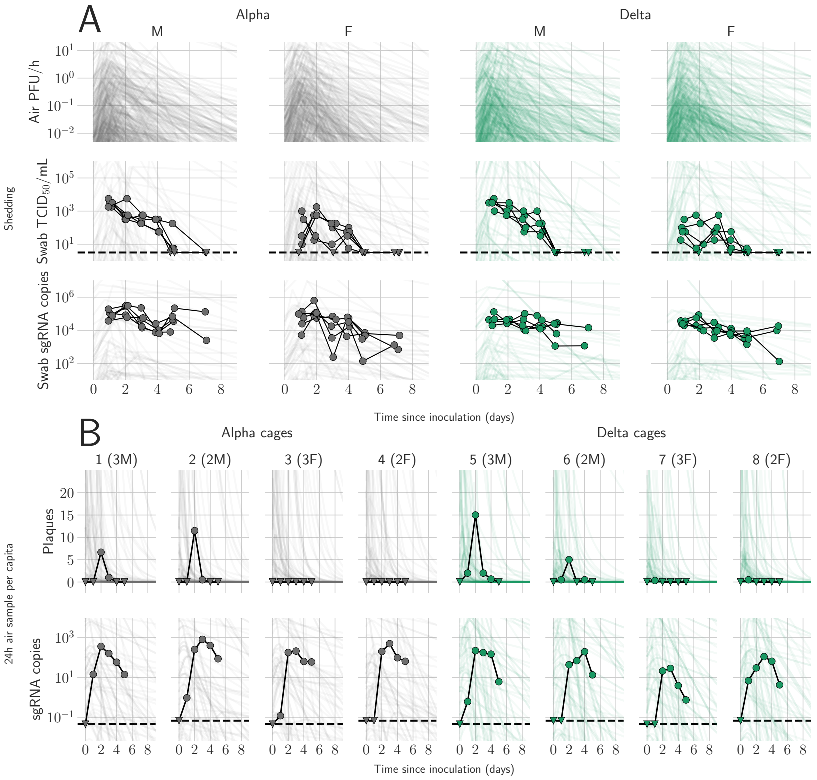

Figure M1: Prior predictive checks. Figure is as main text Figure 1, except that the trajectories showed by the semi-transparent lines are randomly sampled from the prior predictive distribution rather than from the posterior, and 500 lines are drawn in panel A rather than 100, given the greater dispersion relative to the data. Wide range of simulated trajectories relative to the data shows that priors allow for a wide range of a priori plausible kinetics.

#### 5.5 Prior predictive checks

We assessed the appropriateness of prior choices via prior predictive checks. Figure M1 shows a version of main text Figure 1 but where sample trajectories are plotted alongside the data, but drawn from the prior predictive distribution rather than the posterior distribution.

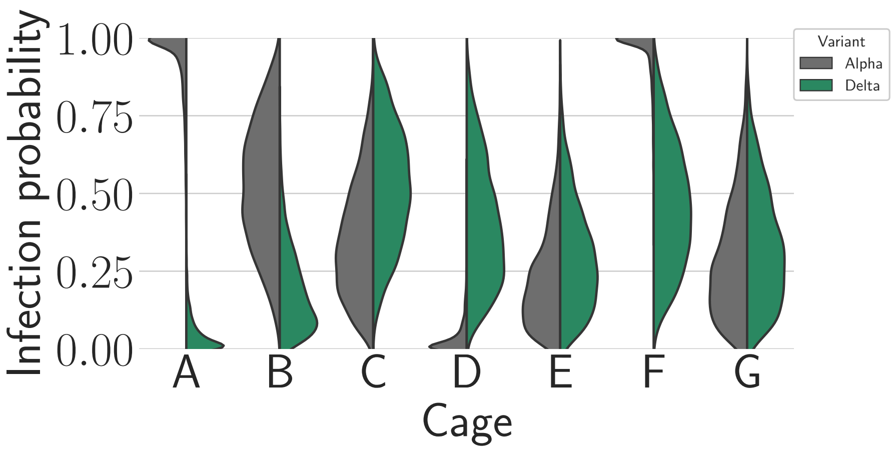

Figure M2: Posterior estimates for the infection probabilities for each variant in each cage. Half-violin plots show posterior densities for Alpha infection probability (gray) compared to Delta infection probability (green). There were very high Alpha infection probabilities in cages A and F.

### 6 Assessing coinfection probabilities

To assess the probability of coinfection, we visualized the infection probabilities for each variant in each cage. Given no interaction between two variants' infection processes, the probability of being coinfected for each hamster $j$ is the product of the hamster's probabilities for each variant:

$$P_{\text{coinf}}(j) = P_{\text{inf}}(1, j) P_{\text{inf}}(2, j) \tag{39}$$

The distribution of the number of coinfections in a given cage or experiment is then the convolution of these individual Bernoulli-distributed outcomes for individuals.

#### 6.1 Results

Figure M2 shows the estimated infection probabilities by variant and cage. Cage F was the only cage in which we observed coinfections, and our model shows that it is indeed the only cage in which both Alpha and Delta clearly had a high probability of causing infections in the sentinels.

We then calculated the posterior estimated probability of coinfection occuring for each hamster in each cage, according to equation 39. Figure M3 shows the resulting estimates.

The model estimates that coinfection probabilities were highest in Cage F simply because both Alpha and Delta infection probabilities were high. In other cages, coinfection probabilities are substantially lower, since at least one variant has a low individual infection probability M2. Cage C (Delta, then Alpha) is the only sequential exposure cage in which the absence of coinfections is perhaps surprising; even there, the data are consistent with a coinfection

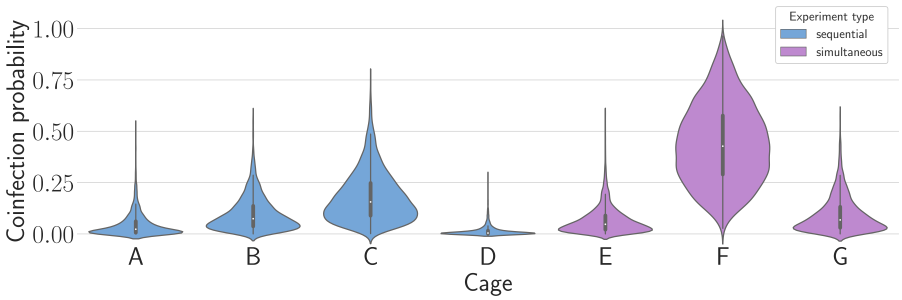

Figure M3: Posterior estimates for coinfection probabilities by cage. Sequential exposures shown in blue, simultaneous exposures shown in pink. Cage F, where coinfections were actually observed, has a substantially higher coinfection probability estimate than other cages.

probability of under 25% or even under 10%, so given that only 5 sentinels were exposed, the absence of a coinfection is consistent with random variation.

Finally, to assess whether the absence of any coinfections in the sequential experiments while several were observed in the simultaneous experiments could be explained by chance, we calculated the posterior distribution for the expected number of coinfections by experiment type (this is the sum of the probability for each cage times the number of sentinels in that cage). The results are shown in Figure M4.

The model suggests that the absence of coinfections in the sequential exposures could simply result from low probabilities in all cages except C; the data are consistent with an expectation of between zero and two coinfections, though more than that would also have been plausible.

Taken together, our results suggest that differences in donor virus dynamics and shedding could readily explain the differences between sequential and simultaneous exposures in our small-N dataset. Identifying or ruling out competitive (or facilitating) interaction among virus variants during sequential versus simultaneous transmission would likely require larger samples.

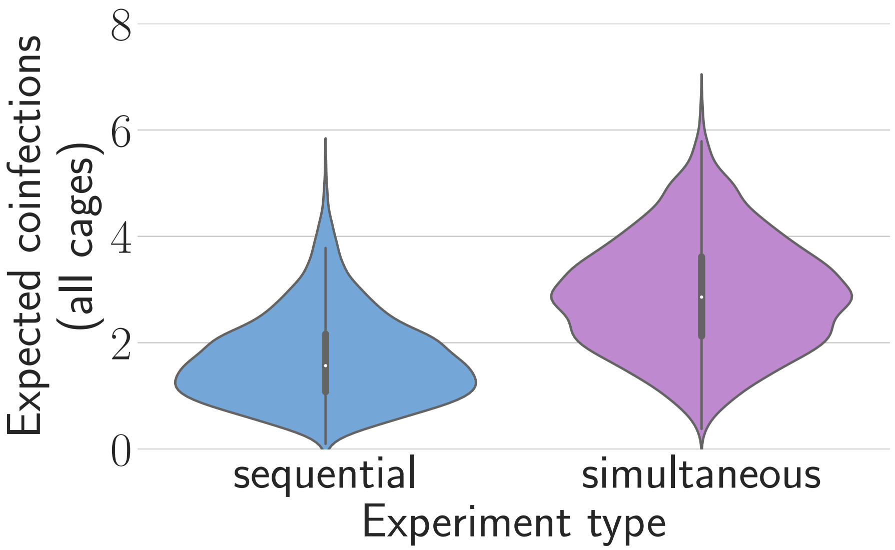

Figure M4: Expected coinfection counts for sequential and simultaneous experiments.

### 7 Computational methods

We implemented and conducted inference from our model in Python using the Numpyro probabilistic programming framework[6]. We drew posterior samples using Numpyro's iterative implementation[6] of the No-U-Turn Sampler (NUTS)[7], a form of Hamiltonian Monte Carlo (HMC). For each model fit, we ran three Markov Chains, each with 1000 warmup steps and 1000 samples. We assessed convergence by examining figures and by confirming sufficient effective sample sizes and the absence of divergent transitions after warmup.

For inference purposes, we set a minimum rate of $10^{-20}$ for all Poisson distributions of "hitting" virions (for filter plaques and virus titration). True 0 rates can cause numerical issue when conducting NUTS sampling with Numpyro. This minimum rate of $10^{-20}$ can be thought of as representing a very small probability of a false positive plaque or well.

We prepared data for modeling and analyzed and visualized output in Python[8]; the packages Numpy[9], Scipy[10], Matplotlib[11], and Polars[12] were particularly critical.

All code and data to reproduce Bayesian inference results, including model fits and model output figures, is available on the project Github repository (https://github.com/dylanhmorris/host-viral-determinants) and archived on Zenodo (https://doi.org/10.5281/zenodo.8396135).

### 8 Additional mathematical details

#### 8.1 Well observation process

Spearman-Karber estimates for a 0.1 mL inoculum give an exact value for the total number of positive wells in a 4 well by 8 serial dilution series (with a an undilute first row $d = 0$, and serial $f$-fold dilutions—here $f = 10$—such that the $d^{\text{th}}$ row has been diluted $f^d$-fold). This allowed us to back-calculate the total number of positive wells. With some additional assumptions, we were then able to calculate the approximate likelihood of observing a given number of positive wells $n$ given a true underlying titer $v$.

We assumed that wells were all positive until some dilution $d$ and wells were all negative or not attempted at dilutions $d + 2$ and higher. That is, only two sequential dilutions are assumed potentially to have a mix of positive and negative wells: $d$ and $d + 1$. This is by far the most probable way to produce a given total number of positive wells, particularly with 10-fold or coarser dilutions, since other hit distributions require some negative wells to occur at substantially higher plated concentrations than some positive wells.

With this assumption made, a given total number of positive wells $n$ has implies a value of this $d$ and a corresponding number $k$ of total positives seen at dilutions $d$ and $d+1$ combined:

$$\begin{aligned} k &= \min\{n, 4 + n \bmod 4\} \\ d &= \frac{n - k}{4} \end{aligned} \tag{40}$$

where mod denotes the modulo operation (remainder when n is divided by 4). Note that while we assume $d$ and $d + 1$ can have a mix of positive wells, we do not assume for certain that they do. When $k = 4$, all positive at $d$ and all negative at $d+1$ is a possible outcome (or, much less probably, all positive at $d+1$ and all negative at $d$); it is just not the *only* possible outcome, as we could also have, e.g., 3 of 4 wells positive at $d$ and 1 of 4 positive at $d+1$.

The approximate log likelihood for observing $n$ positive wells given an underlying virus concentration $v$ is then the sum over the possible ways to generate $k$ positives at dilutions $d$ and $d+1$. Define the random variables $K_d$ and $K_{d+1}$ as the number of wells positive at dilutions $d$ and $d+1$, respectively. Then the probability of observing a certain value of $K_d$ given the underlying virus concentration $v$ is given by a binomial distribution with success probability equal to the single hit probability at dilution $d$, i.e.:

$$P(K_d = k \mid v) = \binom{4}{k} p^k (1-p)^{4-k} \tag{41}$$

For our experiments, $v$ is measured in $\log_{10} \text{TCID}_{50}$ and dilutions are 10-fold, so we have

$$p = 1 - \exp\left(-\log(2) 10^{v-d}\right)$$

This gives us a computable approximate likelihood $\mathcal{L}(n \mid v)$ :

$$\mathcal{L}(n \mid v) \approx \sum_{c=k-4}^{4} \log[P(K_d = c \mid v)] + \log[P(K_{d+1} = k - c \mid v)] \tag{42}$$

#### 8.2 Alternative approach to computing viable virions shed during a time window

This describes a second, equivalent manner of computing $A_v(t_1, t_2)$, the cumulative number of virions shed between $t_1$ and $t_2$ that remain viable at $t_2$. In this instance, we integrate over the time of shedding $t$, but thin the virions shed by considering only the fraction $e^{-\lambda(t_2 - t)}$ that will be viable when sampled at $t_2 \geq t$. This fraction is exact if we treat virion loss of infectivity as exponential at a rate $\lambda$.

$$A_v(t_1, t_2) = m \int_{t_1}^{t_2} V_a(t) e^{-\lambda(t_2 - t)} \, \mathrm{d}t \tag{43}$$

This integral can be computed using the following antiderivative:

$$F(t, a, t_i, t_s) = \int \exp[a(t - t_i) - \lambda(t_s - t)] \, \mathrm{d}t = \frac{1}{a + \lambda} \exp[a(t - t_i) - \lambda(t_s - t)] \tag{44}$$

In our case, $a$ is the exponential growth rate of shedding, with negative $a$ values representing exponential decay, and $t_s \geq t$ is the time of filter removal (so $t_s - t$ is how long a virion deposited at $t$ must retain infectiousness in order to be infectious when the filter is removed).

To compute $A_v(t_1, t_2)$, we have to consider several possible cases. Recall that time is measured relative to the time $t = 0$ that the shedding individuals were inoculated, and $t_a > 0$ is the time of peak air shedding.

If $t_2 < t_a$, the entire sampling period happens before peak air shedding. In that case:

$$A_v(t_1, t_2) = mV_a(t_1)\left[F(t_2, g, t_1, t_2) - F(t_1, g, t_1, t_2)\right] \tag{45}$$

If $t_1 > t_a$, the entire sample is taken after air shedding has peaked. In that case:

$$A_v(t_1, t_2) = mV_a(t_1)\left[F(t_2, -d_{av}, t_1, t_2) - F(t_1, -d_{av}, t_1, t_2)\right] \tag{46}$$

If the air shedding peak occurs during sampling ($t_1 \le t_a \le t_2$), the problem can be solved piecewise:

$$\begin{aligned} A_v(t_1, t_2) = &\int_{t_1}^{t_a} mV_a(t_1)\exp\left[g(t - t_1)\lambda(t_2 - t)\right] \mathrm{d}t + \\ &\int_{t_a}^{t_2} mV_a(t_a)\exp\left[-d_{av}(t - t_a) - \lambda(t_2 - t)\right] \mathrm{d}t \end{aligned} \tag{47}$$

And so applying the antiderivative $F$ from 44:

$$\begin{aligned} A_v(t_1, t_2) = &\, mV_a(t_1)\left[F(t_a, g, t_1, t_2) - F(t_1, g, t_1, t_2)\right] + \\ &\, mV_a(t_a)\left[F(t_2, -d_{av}, t_a, t_2) - F(t_a, -d_{av}, t_a, t_2)\right] \end{aligned} \tag{48}$$

12. Vink, R. *Polars: Blazingly fast DataFrames in Rust, Python & Node.js* https://github.com/pola-rs/polars.
